# Delimiting the cryptic diversity and host preferences of *Sycophila* parasitoid wasps associated with oak galls using phylogenomic data

**DOI:** 10.1101/2022.01.21.477213

**Authors:** Y. Miles Zhang, Sofia I. Sheikh, Anna K.G. Ward, Andrew A. Forbes, Kirsten M. Prior, Graham N. Stone, Michael W. Gates, Scott P. Egan, Linyi Zhang, Charles Davis, Kelly L. Weinersmith, George Melika, Andrea Lucky

## Abstract

Cryptic species diversity is a major challenge for the species-rich community of parasitoids attacking oak gall wasps due to a high degree of sexual dimorphism, morphological plasticity, small size, and poorly known biology. As such, we know very little about the number of species present, nor the evolutionary forces responsible for generating this diversity. One hypothesis is that trait diversity in the gall wasps, including the morphology of the galls they induce, has evolved in response to selection imposed by the parasitoid community, with reciprocal selection driving diversification of the parasitoids. Using a rare, continental-scale data set of *Sycophila* parasitoid wasps reared from 44 species of cynipid galls from 18 species of oak across the US, we combined mitochondrial DNA barcodes, Ultraconserved Elements (UCEs), morphological, and natural history data to delimit putative species. Using these results, we generate the first large-scale assessment of ecological specialization and host association in this species-rich group, with implications for evolutionary ecology and biocontrol. We find most *Sycophila* target specific subsets of available cynipid host galls with similar morphologies, and generally attack larger galls. Our results suggest that parasitoid wasps such as *Sycophila* have adaptations allowing them to exploit particular host trait combinations, while hosts with contrasting traits are resistant to attack. These findings support the tritrophic niche concept for the structuring of plant-herbivore-parasitoid communities.

## 1. Introduction

Tritrophic communities of plants, insect herbivores and associated natural enemies together comprise more than 50% of all described species (Novotny et al., 2010), and include both beneficial ecosystem service providers such as pollinators and biocontrol agents as well as major economic pests of agricultural and forestry. A key aim in evolutionary ecology is to understand the processes that structure this spectacular diversity, and insect-induced galls on plants are veritable cradles of such diversity. Insect galls are highly-structured plant tissues whose development is induced by another organism, and within which the herbivorous immature stages feed on gall tissues and grow to maturity (Price et al., 1987; Rohfritsch & Shorthouse, 1982). An estimated 211,000 species across six insect orders, or ∼4% of estimated global insect species richness, induce galls. Additionally, galls are natural resource-rich microcosms that, in addition to the gall inducer, can support more than 20 species of natural enemies (Askew et al., 2013; Forbes et al., 2016; Weinersmith et al., 2020).

One species rich insect community associated with galls that is well-suited for analysis of tritrophic relationships comprises the North American oak gall wasps (Cynipidae: Cynipini) and their associated hymenopteran natural enemies. The Cynipini induce galls on oaks (*Quercus* spp.) and related Fagaceae, and have a global richness of ∼1000 species in ∼50 genera mostly found in the Northern Hemisphere (Buffington et al., 2020). North America has a relatively high oak species richness (150 species; Cavender-Bares, 2019; Hipp et al., 2018; Manos & Hipp 2021), and an associated high species-richness of oak galling cynipids (∼700 species north of Mexico, Burks 1979). Though scientific study of oak gall system has been an area of active research for well over a century in the Western Palearctic region (e.g. Askew 1961; Bailey et al. 2009; Hayward & Stone, 2005; Nicholls et al. 2017), Nearctic oak gall communities remain relatively poorly known. Though the natural enemies of most North American oak gall wasps remain unknown, the oak galls studied in detail harbor high richness of up to25 species of parasitoids, hyperparasitoids, and inquiline cynipids (herbivorous wasps that are obligate inhabitants of galls induced primarily by other cynipids) (Abe et al., 2007; Forbes et al., 2016; Hayward & Stone, 2005; Schönrogge et al., 1996; Stone et al., 2012; Weinersmith et al., 2020). The parasitoid assemblages attacking regional sets of oak cynipid galls in the Western Palearctic typically overlap, and most of the parasitoids attack multiple host gall types, stimulating ongoing research in the processes that structure cynipid-associated parasitoid communities (Askew et al. 2013; Bailey et al. 2009; Bunnefeld et al 2018).

Oak galls are frequently structurally complex, including characteristic sets of external traits (e.g. spines, hairs, nectar-secreting glands) and internal traits (e.g. internal airspaces, larval chambers that are suspended by radiating fibers or are free-rolling within the gall) (Figure 1), which represent the extended phenotypes of gall wasp genes (Abrahamson & Weis, 1997; Bailey et al., 2009; Hearn et al., 2019; Martinson et al., 2021; Stone & Cook, 1998; Stone & Schönrogge, 2003; Ward et al., 2022). Parasitoid enemies inflict high mortality on cynipid gall inducers, and the Enemy Hypothesis posits that these gall structural traits have likely evolved as defenses against natural enemies, which then drives reciprocal phenotypic evolution in relevant traits of parasitoid wasps, such as ovipositor lengths (Bailey et al., 2009; Price et al., 1987; Stone & Schönrogge, 2003). The complexity of this system is further enriched by the cyclical parthenogenic life cycles of most Cynipini, with obligate alternation between spring sexual and autumn asexual generations that induce morphologically distinct galls (often on different parts of the tree), which host different sets of natural enemies (Bailey et al., 2009; Stone & Schönrogge, 2003). A general property of Western Palearctic cynipid communities is that most of the parasitoids involved attack multiple hosts, with some attacking over 100 gall types (Askew et al. 2013). The extent to which this is true of parasitoids in other global oak cynipid communities is unknown, but it is central to understanding the relationship between gall traits and parasitoid phenotypic evolution (Hayward & Stone 2005, Bailey et al. 2009). The general hypothesis is that where gall wasps show high diversity in relevant gall traits, these will influence associated parasitoid assemblages.

**Figure 1.**
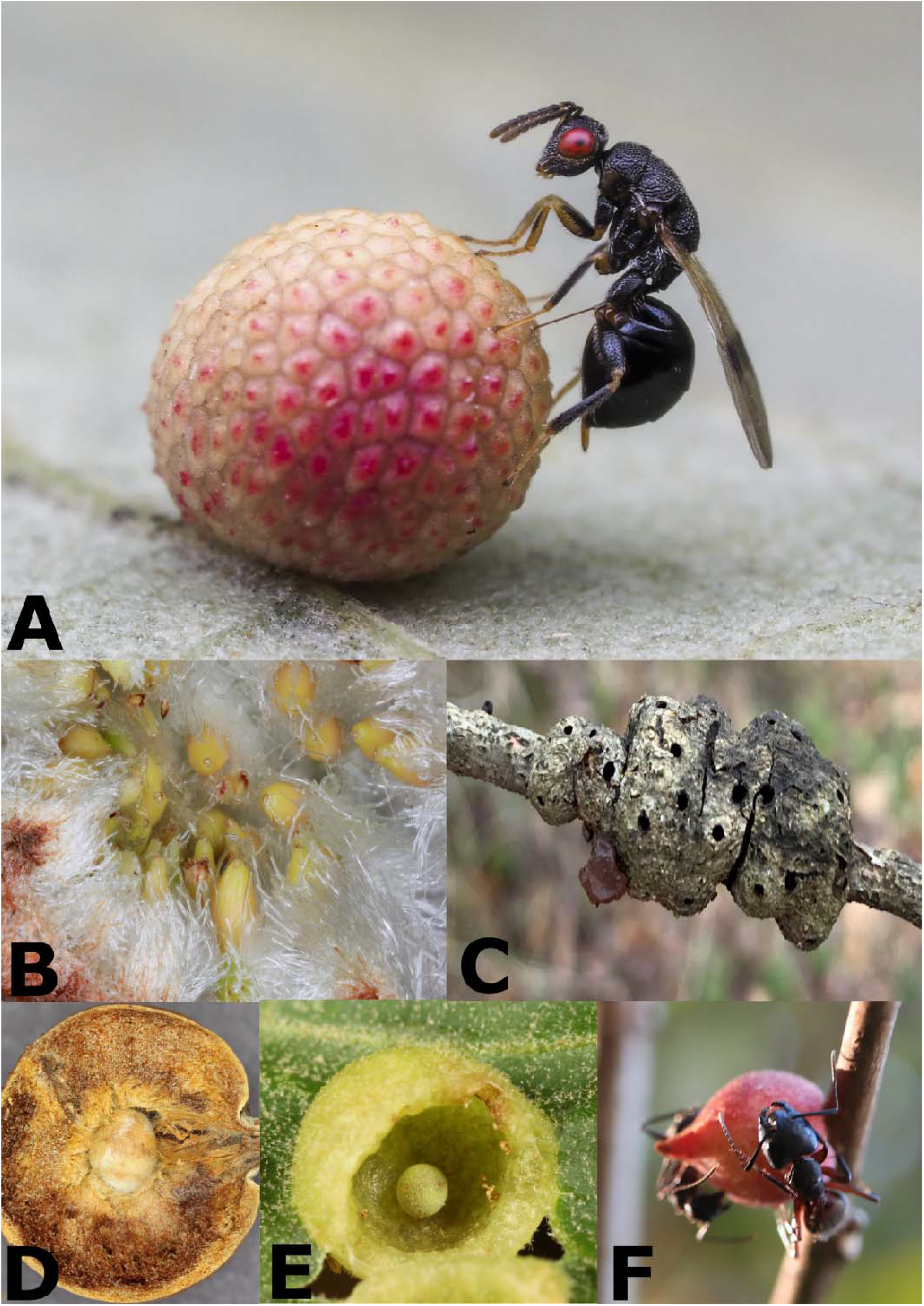
**A**. *Sycophila* sp. ovipositing into a detachable leaf gall of *Acraspis pezomachioides*. **B**. Wooly bud gall of *Callirhytis seminator* with larval chambers. **C**. Integral stem gall of *Callirhytis quercuspuncatata* with exit holes. **D**. Cross-section of woody stem gall of *Disholcaspis quercusglobulus*. **E**. Free-rolling larval chamber of integral leaf gall of *Dryocosmus quercuspalustris*. **F**. *Camponotus* ants feeding on nectar secreted by the gall of *Disholcaspis quercusmamma*. Photo A by Carroll Perkins, B/D by Anna Ward, C/F by Jeff Clark, D by Charley Eiseman.

One such associated group of parasitoids is the genus *Sycophila* (Hymenoptera: Eurytomidae, Figure 1A). *Sycophila* are cosmopolitan in their distribution, and are primarily endoparasitoids of endophytic insects including gall inducers (Askew et al. 2006; 2013; Balduf 1932; Gómez et al., 2013), although a recently described parasitoid species is thought to be an ectoparasitoid of a eulophid gall on *Smilax* (Gates et al. 2020). *Sycophila* are often identified based on subtle differences in adult coloration, host species, and/or geographic distribution (Balduf, 1932; Claridge, 1959). Some described North America species have a wide host repertoire (e.g. *Sycophila quercilanae* = 19 gall types on various oak species, *Sycophila occidentalis* = 12, *Sycophila varians* = 11, *Sycophila dorsalis* = 9), though lack of detailed study suggests that these host repertoires are likely underestimates, and they are low compared to some Western Palearctic species in oak galls (e.g. *Sycophila biguttata* = 80, *S. variegata* = 41) (Askew et al. 2013; Balduf, 1932; Noyes, 2020). The high level of sexual dimorphism, morphological plasticity, and poorly known biology have further confounded species delimitation within this group (Davis et al., 2018; Gómez et al., 2013; Li et al., 2010; Lotfalizadeh et al., 2008; Smith-Freedman et al., 2019). Inability to reliably identify *Sycophila* greatly hampers our understanding of the ecology and evolutionary history of their interactions with host galls, and also limits our understanding of *Sycophila* serving as potential biocontrol agents of pestiferous gall wasps such as the invasive Asian chestnut gall wasp, *Dryocosmus kuriphilus* (Dorado et al. 2020), or the North American species *Zapatella davisae, which* damages black oaks in the New England area (Davis et al., 2018, Smith-Freedman et al., 2019).

Assessing species richness and revealing axes of host specialization for gall parasitoids requires a well-resolved and stable parasitoid taxonomy, which in turn requires an integrative approach. Single or multi-locus approaches for taxon delimitation that use genes such as the mitochondrial loci COI and Cytb have been used extensively to understand gall community diversity (Ács et al., 2010; Davis et al., 2018; Forbes et al., 2016; Gil-Tapetado et al., 2021; Kaartinen et al., 2010; MacEwen et al., 2020; Nicholls et al., 2018; Nicholls et al., 2010; 2018; Sheikh et al., 2022; Smith-Freedman et al., 2019; Ward et al., 2020; Weinersmith et al., 2020; Zhang Y.M. et al., 2014; 2019a), often in combination with morphological and/or ecological data. However, results from one or a few genes can be limited in resolution, and single locus mitochondrial COI barcodes are known to be misleading due to confounding factors such as incomplete lineage sorting and introgression within both gall wasps and their associated parasitoids (Nicholls et al., 2012; Rokas et al., 2003; Zhang Y.M. et al., 2021). Nevertheless, molecular approaches remain attractive due to the demonstration of morphologically cryptic species in many parasitoid taxa with wide host repertoires, including members of oak gall wasp communities (Kaartinen et al. 2010; Nicholls et al. 2010, 2018). Rapid development and increased availability of tools designed to capture genomic DNA has led to their increase use in studies of phylogenomics, biogeography, demography, host shifts, and tritrophic interactions of gall communities (Blaimer et al., 2020; Brandão-Dias et al., 2022, Bunnefeld et al., 2018; Driscoe et al., 2019; Samacá-Sáenz et al., 2020; Walton et al., 2021; Ward et al., 2022; Zhang Y.M. et al., 2020). Additionally, targeted capture methods such as Ultraconserved Elements (UCEs, Faircloth et al., 2012, reviewed in Zhang Y.M. et al., 2019b) have been shown to be complementary or superior to DNA barcodes for resolution of deep phylogenetic relationships and species delimitation in Hymenoptera, and can be amplified even from older museum samples (Branstetter & Longino, 2019; Gueuning et al., 2020; Ješovnik et al., 2017; Longino & Branstetter, 2021; Prebus, 2021; Samacá-Sáenz et al., 2020). Thus UCEs are an appealing approach with which to validate species status in morphologically challenging taxa such as *Sycophila*.

The goals of this study are two-fold: **1)** To delimit – using molecular, ecological, and morphological data – putative species among a representative collection of *Sycophila* reared from galls of 44 species of oak gall wasps from 18 oak tree species across the USA. Our sampling targeted two axes known to structure gall wasp-parasitoid associations: different gall wasp faunistic zones *sensu* Weld (Hayward & Stone 2006), and hosts on different oak sections (Bailey et al. 2009). **2)** To further delimit axes of adaptation (∼ host repertoire) and evolutionary histories of host use of North American *Sycophila*. We hypothesize that if parasitoids have cospeciated with their gall wasp hosts, and gall trait combinations are phylogenetically conserved, then closely related *Sycophila* taxa should attack galls with similar traits. Alternatively, if parasitoid associations are structured by host gall traits, then structurally similar galls may be attacked by phylogenetically diverse (ie closely-related and distantly-related) parasitoid lineages. Whether phenotypically similar host galls are closely related or not depends on the phylogenetic pattern of host gall trait evolution: if host gall traits have evolved convergently in North America (as they are known to in the Western Palearctic; Stone and Cook 1998; Cook et al. 2002), we expect a single parasitoid to attack a set of unrelated but phenotypically similar host galls.

## 2. Materials & Methods

### 2.1 Taxon Sampling

The *Sycophila* specimens used in this study were collected through long term collaborative research on the North American oak gall fauna by the authors and their respective collaborators, including many students. In brief, mature galls were collected and the inhabitants were reared from individual galls or from mass rearings of a single gall type (Table S1, Fig. S2). For each gall we recorded the host plant and we scored gall external and internal morphological traits (Table S2) using mature galls from collections at the Smithsonian National Museum of Natural History, published terminologies from www.gallformers.org and existing literature (Deans et al., 2021; Weld, 1959). The trait set includes discrete binary or categorical characters describing gall position on plant (acorn, catkin, leaf, petiole, stem), attachment type (integral, detachable), external morphology (smooth, textured, leaf bract, sticky, spiny, wooly), and internal morphology (woody, hollow, fleshy, free-rolling, or radiating fiber). For these traits, integral attachment refers to galls whose tissues are broadly continuous with plant organs, such that the gall does not typically detach or dehisce from the plant when mature. External morphological traits include surface texture (for which the textured state indicates uneven surfaces that can be knobbled or rugose, Fig. 1A), and presence/absence of other traits (ant-recruiting nectaries, coatings of spines or wool, Fig. 1B, 1F) implicated in defense against parasitoids in other studies (Bailey et al. 2009; Nicholls et al. 2018). Internal morphological traits include the texture of gall tissues (woody, hollow, fleshy) and two internal traits (free rolling larval chamber and a larval chamber suspended in the center of the gall by fine radiating fibers, Fig. 1E) also associated with reduction in successful parasitoid attack (Bailey et al. 2009; Martinson et al. 2021). We scored mature gall size as a categorical variable, with 1 representing large (2–15 cm) galls, 2 as medium (0.5–2 cm) galls, and 3 as small (<0.5 cm) galls (Table S2). As gall hardness varies substantially with gall age, we did not include this trait in the current study in order to standardize traits across different collectors/events. We categorized gall wasp distributions using the biogeographic regions Pacific Slopes, Southwest, and Eastern United States established by Weld (1957, 1959, 1960).

Host trees were scored based on sections in *Quercus* sensu Manos & Hipp (2021): *Lobatae* (red oaks), *Protobalanus* (intermediate or golden cup oaks), *Quercus s*.*s*. (white oaks), and *Virentes* (live oaks). Adult *Sycophila* specimens were identified to species morphologically whenever possible using Balduf (1932) and double-checked with the type specimens at the US National Museum of Natural History (NMNH), Washington D.C. Where morphology-based identities could not be confidently assigned, we identified *Sycophila* specimens based on a combination of wing band, body coloration (e.g. *Sycophila* sp1–7, Fig. S1), and host information in cases where no matches were found. Detailed information on each of the *Sycophila* species are provided in Supplementary Figure 1. Representatives from each of the morphospecies were selected for downstream molecular analyses. One to several representatives of each *Sycophila* morphospecies from each different host gall type and/or widely separated locations were sequenced to sample the greatest possible degree of genetic variation based on host, geographic distance, and morphological variation. Secondary voucher specimens from the same collection events as the samples destructively sampled for DNA extraction are deposited at the NMNH and University of Iowa when possible, but some morphologically cryptic singleton species were only discovered after sequencing and thus do not have morphological vouchers. Habitus images were obtained using a Macropod imaging system consisting of a Canon EOS 5D Mark II digital SLR camera with a 65mm macro lens, illuminated with a Dynalite MP8 power pack and lights. Images were captured using Visionary Digital proprietary software as TIF with the RAW conversion occurring in Canon Digital Photo Professional software. Image stacks were mounted with Helicon Focus 6.2.2. Images were edited in Adobe Photoshop.

### 2.2 DNA Extractions, COI sequencing

Due to the small size (<3mm on average) and low DNA yield (∼1ng/μL), representative specimens of *Sycophila* (n = 89) were destructively sampled at either the University of Iowa or Rice University TX, USA. One third of specimens were extracted using the DNeasy Blood and Tissue Kit (Qiagen, Valencia, CA, USA), while later extractions used a CTAB/PCI extraction approach (Chen et al., 2010) as it yielded higher quality and quantity of DNA. Approximately 650bp of COI was amplified using either COI_pF2: 5′ ACC WGT AAT RAT AGG DGG DTT TGG DAA 3′ and COI_2437d: 5′ GCT ART CAT CTA AAW AYT TTA ATW CCW G 3′ primers (Kaartinen et al., 2010), or, for most of the specimens, with an in-house forward primer Syco_2: 5’-TTC CWG ATA TRG CTT TYC C -3’ and COI_2437d. The Syco_2 primer was designed to reduce degeneracy while still overlapping with the COI region amplified using the Kaartinen et al. (2010) primers. Forward and reverse Sanger sequencing was done on an ABM 3720 DNA Analyzer (Applied Biosystems, Foster City, CA) in the University of Iowa’s Roy J. Carver Center for Genomics, and reads were processed in Geneious v8 (Biomatters Inc., San Diego, CA) for final consensus sequences. Additional COI sequences of *Sycophila* reared from asexual generation of *Zapatella davisae* (Smith-Freedman et al., 2019) and *Belonocnema kinseyi* (Forbes et al. 2016) were downloaded from GenBank for a total of 165 sequences, along with the sequence of *Eurytoma longavena* which was used as an outgroup (Zhang Y.M. et al., 2014).

### 2.3 UCE Data Collection

The UCE pipeline was conducted in the Laboratories of Analytical Biology (LAB) at the Smithsonian Institution’s National Museum of Natural History (NMNH, Washington, DC, USA). The protocol largely follows the standard pipeline for capturing and enriching UCE loci from Hymenoptera (Branstetter et al., 2017; Zhang Y.M. et al., 2019b). Briefly, the DNA extracts from 30 destructively-sampled individuals using the DNeasy Blood and Tissue Kit (Table S1) were chosen based on high DNA quality, and the Kapa Hyper Prep library preparation kit (Kapa Biosystems Inc., Wilmington, MA, USA) was used along with TruSeq universal adapter stubs and 8-bp dual indexes (Glenn et al., 2019), combined with sheared genomic DNA and amplified using PCR. We followed the myBaits probes V4 protocol (ArborBiosciences, Ann Arbor, MI, USA) for target enrichment of the pooled DNA libraries but instead used a 1:4 (baits:water) dilution of the custom Hymenoptera 2.5Kv2P developed by Branstetter et al. (2017) at 65ºC for 24 hours. The combined library was sequenced on Illumina NovaSeq 6000 (150-bp paired-end, Illumina Inc., San Diego, CA, USA) at Novogene Corporation Inc. (Sacramento, CA, USA).

### 2.4 UCE Data Processing and Alignment

We used the PHYLUCE v1.6.8 pipeline (Faircloth, 2015) to process UCE data. Adapters were trimmed using Illumiprocessor and Trimmomatic (Bolger et al., 2014; Faircloth, 2013), and assembled using SPAdes v3.14.0 (Bankevich et al., 2012). The assemblies were aligned using MAFFT v7.490 (Katoh & Toh, 2008), and trimmed using Gblocks (Castresana, 2000) using the following settings: b1=0.5, b2=0.5, b3=12, b4=7. Additionally, we used Spruceup v2020.2.19 95% lognormal distribution or manual cutoff of select samples to remove any potentially misaligned regions as they can produce exaggerated branch lengths (Borowiec, 2019). We selected the 50% complete matrix with 1456 loci that are present in ≥50% of the taxa (15/30) as the final dataset. A 75% matrix (627 loci) was also tested to ensure topological consistency with a small set of more data-complete specimens. The topology of this tree was entirely concordant (data not shown). Phylogenetic summary statistics were calculated using AMAS v0.98 (Borowiec, 2016). Additionally, fragments of mitochondrial DNA COI were extracted from the UCE contigs using PHYLUCE script phyluce_assembly_match_contigs_to_barcodesto be used in conjunction with full COI barcodes whenever possible.

### 2.5 Phylogenetic Analyses

We conducted phylogenetic analyses under the maximum likelihood (ML) criterion with IQ-TREE v2.03 (Minh et al., 2020) for the COI data, using the best model (GTR+F+I+G4) chosen by ModelFinder (Kalyaanamoorthy et al., 2017), and 1000 ultrafast bootstrap replicates for nodal support (UFB, Hoang et al., 2017).

The UCE data were also analyzed using the ML criterion with IQ-TREE, using partitions based on Sliding-Window Site Characteristics of Site Entropy (SWSC-EN, Tagliacollo & Lanfear, 2018)), and partitioned using the rcluster algorithm in PartitionFinder2 via RAxML using default settings (Lanfear et al., 2014, 2016; Stamatakis, 2006). To assess nodal support, we performed 1000 UFB, along with “-bnni” to reduce risk of overestimating branch supports; and a Shimodaira-Hasegawa approximate likelihood-rate test (SH-aLRT, Guindon et al., 2010) with 1000 replicates. Only nodes with support values of UFB ≥ 95 and SH-aLRT ≥80 were considered robust.

### 2.6 Delimitation of Putative Species

We used multiple molecular species delimitation methods in combination with geographical, ecological, and morphological data to delimit putative *Sycophila* species. Because many collections were made from different galls and host trees in the same geographic locations, correspondence between genetic differences, wing pattern differences, and different host associations provides strong indirect support for limited gene flow between sympatric individuals. A complete discussion of each putative *Sycophila* species, including representative body and wing images for most species, is provided in the supplemental materials, Fig. S1.

For the COI data, we explored three popular molecular species delimitation methods: 1) Assemble Species by Automatic Partitioning (ASAP, https://bioinfo.mnhn.fr/abi/public/asap/, Puillandre et al., 2021), an extension of the Automatic Barcode Gap Discovery method (ABGD, Puillandre et al., 2012) was performed using the default setting using uncorrected p distance. 2) The Bayesian Poisson Tree Processes (bPTP, https://species.h-its.org/, Zhang J.et al., 2013) was performed on the same dataset using the default settings of 200,000 MCMC generations, thinning of 100, and 0.1 burn-in. 3) The Generalized Mixed Yule Coalescent (GMYC, https://species.h-its.org/gmyc/, Pons et al., 2006) was performed on an ultrametric input tree generated in BEAST2 v. 2.2.7 (Bouckaert et al., 2014). The JC69 substitution model and a strict molecular clock with a fixed rate of 1.0 were used, following a Yule model with a uniform distribution for “birthRate”. The analysis ran for 10 million generations, with sampling every 1,000 generations. Convergence was confirmed with ESS above 200 in all categories using Tracer v1.7 (Rambaut et al., 2018). The resulting tree was analyzed using the single-threshold version of Splits R package (Ezard et al., 2009). Intra- and interspecific divergence among the species were calculated using MEGA11 (Tamura et al. 2021) using uncorrected distance.

For the UCE data, we also performed three species delimitation methods under the multi-species coalescent model (MSC). 1) We tested the full UCE dataset using SODA v1.0 (Rabiee & Mirarab, 2020), which delimits species boundaries using quartet frequencies. Gene trees were generated using the best models selected from ModelFinder, and SODA was performed without using a guide tree. 2) We performed allelic phasing on the UCE loci following Tutorial II of the PHYLUCE pipeline, which has been shown to improve species delimitation (Andermann et al., 2019). The reads were aligned to the assembled contigs, and the data were re-aligned and trimmed before single nucleotide polymorphisms (SNPs) were extracted from the phased UCE loci using SNP-sites V2.5.1 (Page et al., 2016), selecting one random SNP per locus to avoid linkage disequilibrium. The phased SNPs were analyzed using STACEY (Jones, 2017) as implemented in BEAST2 with model selection performed for each locus using the bModeltest option and corrected for ascertainment bias (Bouckaert & Drummond, 2017). Species trees were estimated using a strict clock at 1.0 under the Fossilized Birth Death model (Heath et al., 2014), using a value of 1 × 10^−4^ for the collapseHeight parameter, bdcGrowthRate = log-normal (M = 4.6, S = 2); collapseWeight = beta (alpha = 2, beta = 2); popPriorScale = log-normal (M = −7, S = 2); relativeDeathRate = uniform (upper = 1.0). The analysis ran for 10 million generations, sampling trees every 100,000 generations. Convergence was confirmed with ESS above 200 in all categories using Tracer v1.7 (Rambaut et al., 2018), and the sampled species trees were visualized with DensiTree 2.2.7 (Bouckaert, 2010). 3) We selected a subset of 50 phased UCE loci with the greatest number of parsimony-informative sites to reduce computational time using the Phyloch R package (Heibl, 2008). We then used BPP v4.3.8 (Yang, 2015) with tau and theta parameters estimated using the A00 analysis on the fixed SWSC tree, without delimitation. Using the resulting parameters, we then performed the rjMCMC species delimitation algorithm A01 (species delimitation = 1 1 2 1), with the number of MCMC generations to 300 K, sampling every five generations, with a 25% burn in.

### 2.7 Principal Coordinates Analysis of Gall Traits

To ascertain whether groups of *Sycophila* species attack gall wasp species with particular gall morphology, we scored each gall for defensive morphological traits (scored as 1 if present) and gall size (large gall size of 2–15cm scored as 1 as putative defense). We then calculated Gower’s dissimilarity, which is appropriate for a mix of binary and categorical variables, between all gall pairs on a gall wasp species × trait matrix (Laliberte & Legendre 2010). We then performed a principal coordinates analysis (PCoA) and projected gall wasp species in trait space by creating a biplot with PCoA1 and PCoA2, and plotted loadings representing gall traits (Fig. S5) (Dehling et al. 2015). Next, to project *Sycophila* species in interacting gall wasp species trait space (i.e., “interaction” trait space), for each *Sycophila* species (Table S5), we calculated the interaction centroid as the center of gall wasp species that each *Sycophila* species interacts with. We plotted the centroids for each *Sycophila* species in biplots, to visualize if closely or distantly related *Sycophila* assemble on galls with certain sets of traits.. We used R v4.1.1 (R Core Team 2021) and the following R packages, ‘labdsv’, ‘vegan’, and ‘ape’, to perform analyses and make biplots (Roberts 2019, Oksanen et al. 2020, Paradis et al. 2021).

Next, to test if distances in interaction trait space between paired *Sycophila* species are correlated with phylogenetic distances, we created a matrix of Euclidean distances between each pair of *Sycophila* species in interaction trait space (smaller values meaning *Sycophila* species are attacking galls with similar defensive traits). We also created a matrix of pairwise interspecific genetic distances (a proxy for relatedness among *Sycophila* species) using the uncorrected distance using default settings in MEGA 11 (Tamura et al. 2021). We then performed a Mantel test between the evolutionary distance matrix and the interaction trait distance matrix in R using in the package ‘vegan’ (Oksanen et al. 2020). If closely related species of *Sycophila* cluster together in gall trait space, it would suggest they are attacking structurally similar galls. Conversely, if distantly related species of *Sycophila* are attacking galls with similar traits, this would suggest they are convergently targeting specific gall traits to overcome. However, without knowing whether the gall structures have evolved convergently (i.e. phylogeny of the galler), we cannot tease apart whether or not *Sycophila* have cospeciated with their hosts. Mantel tests are used to study relationships among dissimilarities in dissimilarity matrices (Legendre et al. 2015), or in our case whether related *Sycophila* attack structurally similar galls. Recent papers have raised concerns about the power of the Mantel test in specific contexts (Harmon & Glor 2010, Legendre & Fortin 2010, Guillot & Rousset 2013). One concern is the inflation of Type I error, including for dissimilarity matrices of hierarchical phylogenetic distances (Harmon & Glor 2010). Since we find no relationship between genetic distance and interaction trait distances this issue is not a concern in the interpretation of our results. Despite the controversy of Mantel tests in certain contexts, they are still used to compare genetic and trait distances among populations and species (e.g. Borcard & Legendre 2011, Schwallier et al. 2015).

## 3. Results

### 3.1 COI Data

The final COI dataset consisted of 165 *Sycophila* specimens, reared from 44 different oak gall wasp species (27 asexual generation, 17 sexual generation) collected on 18 different oak species (Table S1, Figure S2). Most sequence lengths were 655bp, except for the *Z. davisae* parasitoids from GenBank, which were 414bp due to primer differences, and barcode slices from the UCE contigs which ranged from 193–655bp.

The three species delimitation programs, ASAP, bPTP, and GMYC, delimited 35 (7 sequences removed due to not overlapping), 42, and 40 putative species (Figure 2), respectively. In instances where sequence-based delimitation methods disagreed (*S. nr. foliatae2, S. nr. flava, S. nr. globuli, S. globuli*), we used the most conservative estimate, reducing the final number of putative species down to 35 (Figure 2, S1). It is worth noting that some putative species have low bootstrap support (e.g. *S*. nr. *foliatae*-1 and *S. globuli*, Fig. S3), or could represent population level genetic differences without clear host or geographical differences (e.g. *S*. sp1, *S*. sp2, Fig. 2). The intraspecific divergence ranged from 0.0–4.0% (Table S3), while the interspecific divergence ranged from 4.2–17.3% (Table S4). We assigned the putative species into six species groups based on morphological and genetic similarities (Figure 2).

**Figure 2.**
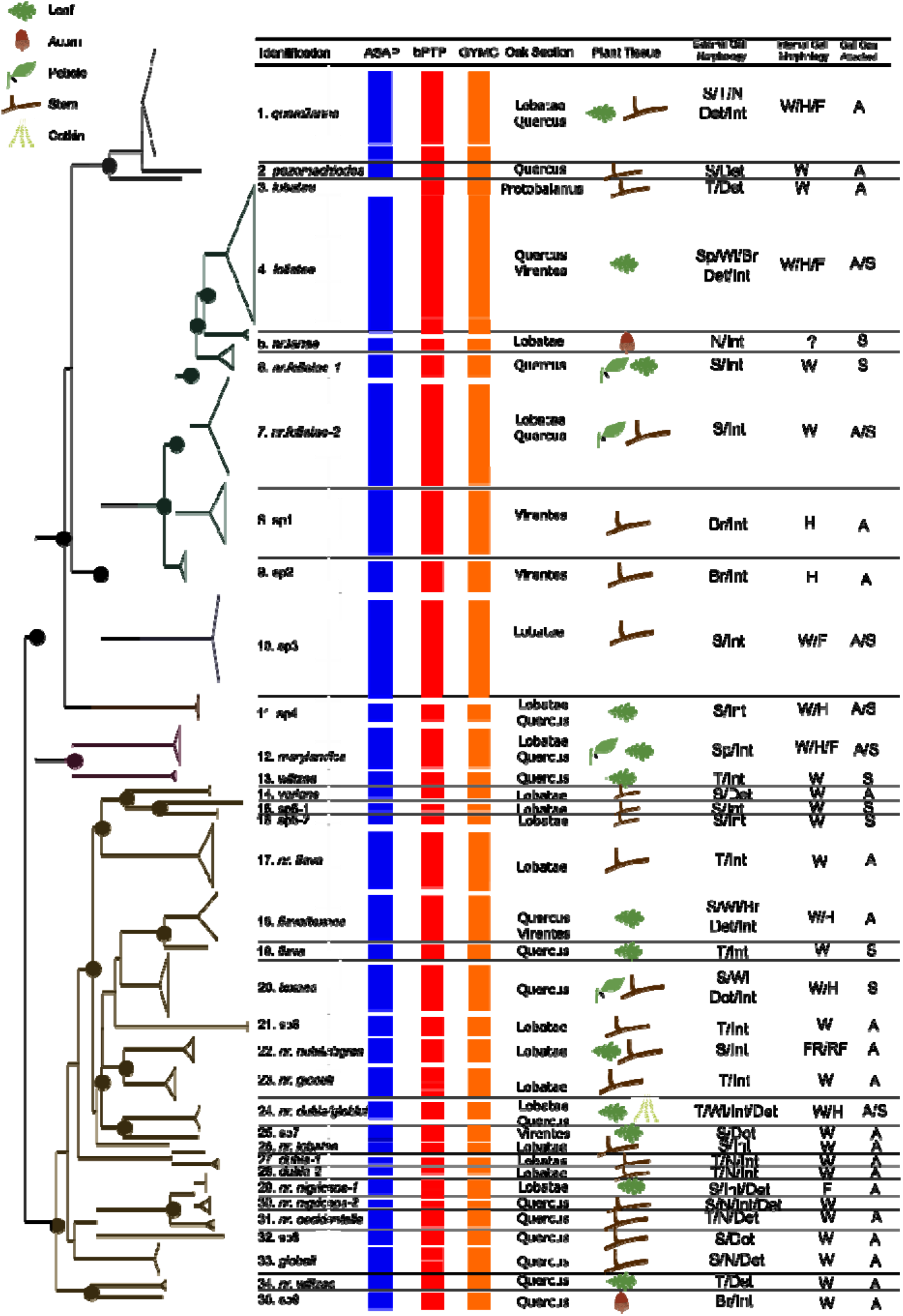
Overview of all COI data used in inferring *Sycophila* diversity associated with North American oak galls. Left: simplified COI phylogeny of *Sycophila* included in this study (see Supp Fig. S3 for full tree). ‘Identification’ describes putative species assignments based on the sum of information to the right of this column. ASAP, bPTP, and GYMC columns indicate assignments of individuals into groups by these respective algorithms. ‘Oak Section’, ‘Plant Tissue’, ‘External Gall Morphology’, ‘Internal Gall Morphology’ and ‘Gall Generation Attacked’ refer to ecological characters for *Sycophila* in each clade, and example photos of galls are shown in Fig. S2. Abbreviations as follows: External Gall Morphology: Br = Leaf Bract, S = Smooth, N = Nectar, T = Textured, Sp = Spines, Wl = Wool, Int = Integral, Det = Detachable; Internal Gall Morphology: W = Woody, H = Hollow, F = Fleshy, FR = Free-Rolling, RF = Radiating Fiber. Black dots represent bootstrap support with ≥75% support. Colored clades correspond to species groups.

Host richness of the putative 35 *Sycophila* species ranged from 1–12 gall wasp species, and we categorized *Sycophila* as 24 extreme specialists (1 host), 10 specialists (2–11 hosts), and one generalist (11+ hosts) following Bailey et al. (2009) (Table 1). *Sycophila quercilanae* had the broadest host repertoire, being reared from 12 gall species from eight different tree species. Two species of *Sycophila* (*S. dubia, S. globuli*) with more than one wasp host species were restricted to hosts from the same genus, while others such as *S. quercilanae* are recorded from hosts in eight different wasp host genera. In terms of the traits of gall wasps attacked by individual putative *Sycophila* species, 2–5 external and 1–3 internal gall morphological traits are observed, found on 1–2 different plant tissues. Most putative *Sycophila* species were reared exclusively from asexual generation galls (20/32), while the remaining species were either reared exclusively from sexual generation galls (6 species), or were reared from both sexual and asexual generations (6 species; Fig. 2). Tree association for *Sycophila* species ranged from 1–7 oak tree species, reflecting either one or two *Quercus* Sections. Most putative species (28/35) were reared from galls on only one *Quercus* Section, while two species (Fig. 2, S1, *S. foliatae, S. flava/texana*) were associated with Sections *Quercus s*.*s*. and *Virentes*, which together form a monophyletic group within the subgenus *Quercus s*.*l*. (Hipp et al., 2018), and six species were associated with both Sections *Quercus* and *Lobatae*.

**Table 1.**
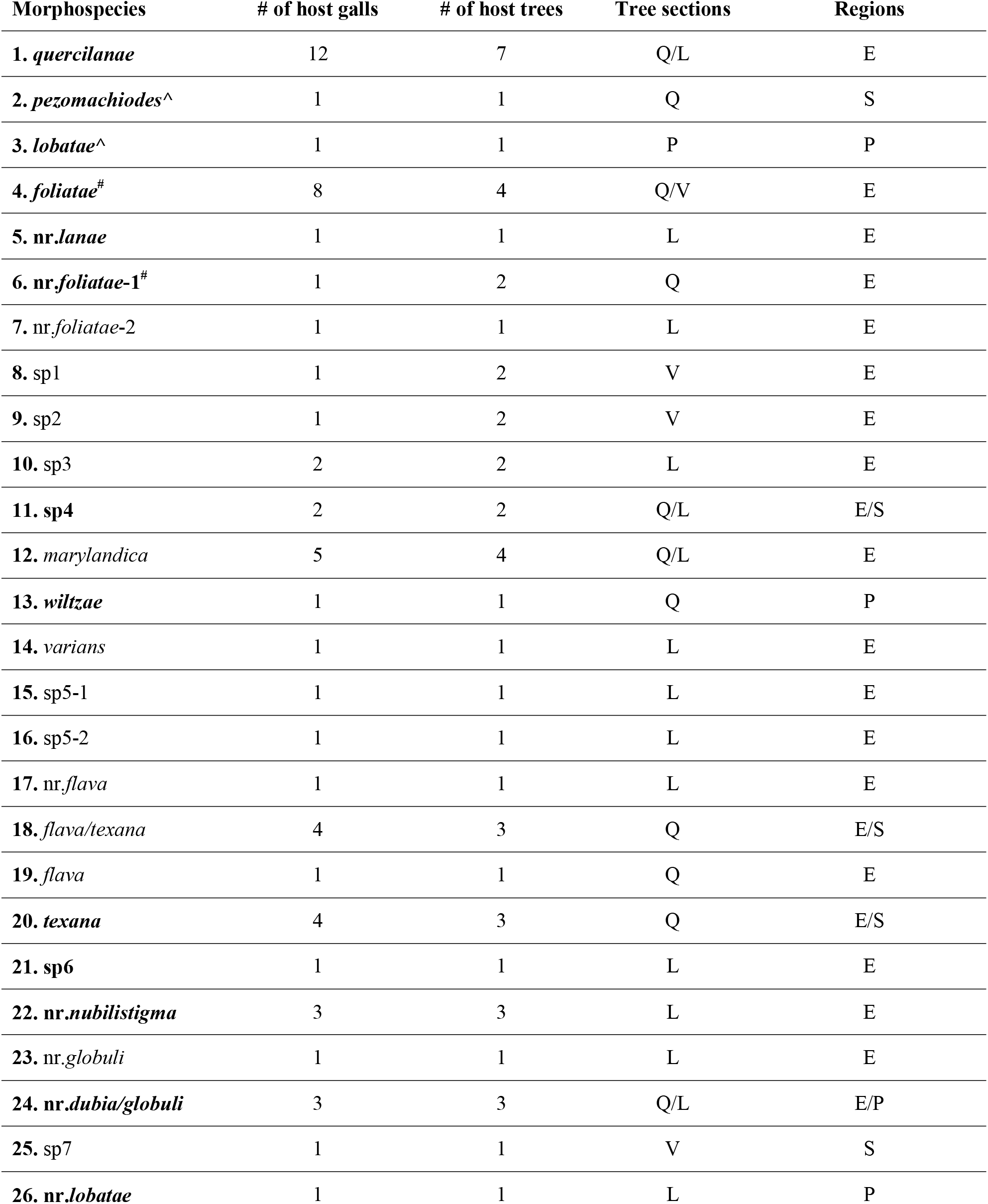
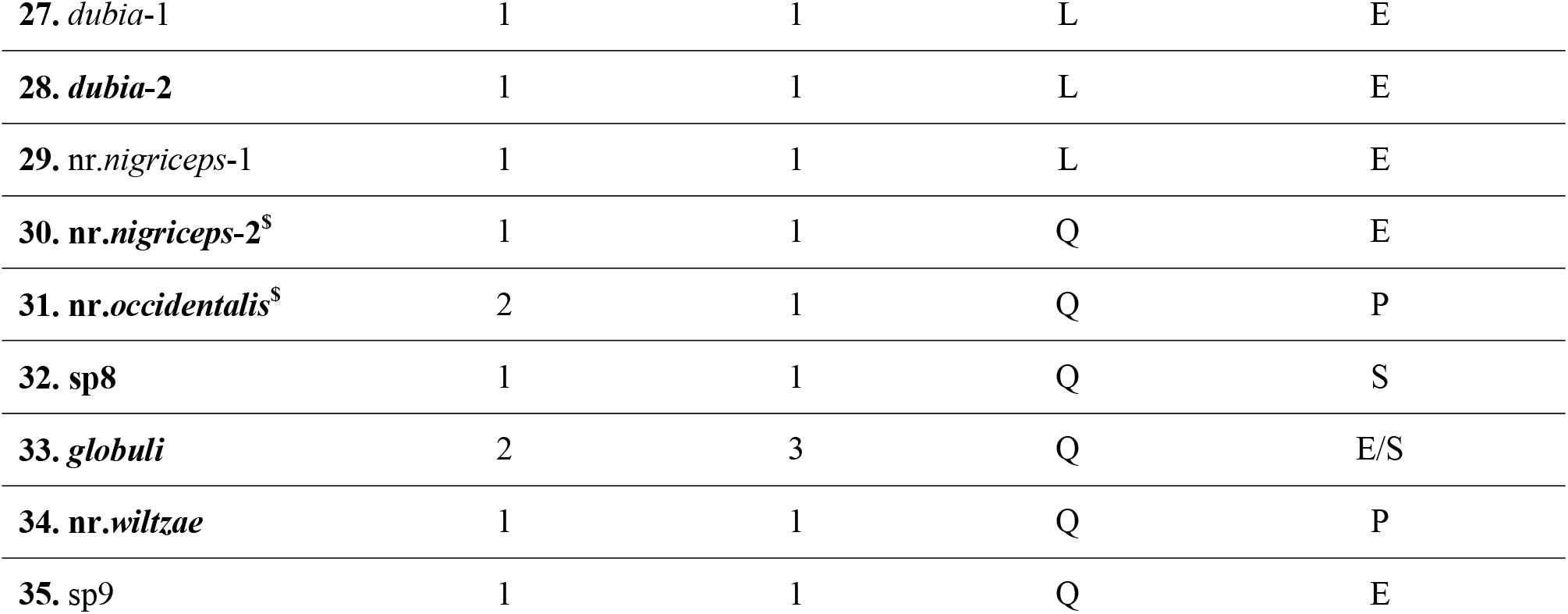
Summary of *Sycophila* COI morphospecies and their host range; bold indicates clades with COI and UCE data. Symbols ($#^) indicate the clades are grouped together in UCE data. Tree Sections Q = *Quercus s*.*s*., P = *Protobalanus*, L = *Lobatae*, V = *Virentes*; Regions E = Eastern US; S = Southwestern US, P = Pacific Slope.

### 3.2 UCE Data

The concatenated UCE 50% matrix was 588,371 bp long, after removing the 0.95 lognormal cutoffs using AMAS, the final matrix is 113,221 variable sites (19.2%) and 34,782 parsimony informative sites (5.9%), with 41.2% missing data.

Species delimitations using UCE data are largely congruent with COI-based results for 19 of the 35 morphospecies, where both data types are available. UCE data supported 22 putative species using the unphased data in SODA and with phased SNPs of the top 50 most parsimoniously informative loci in BPP, while STACEY identified 17 species using the full set of phased SNPs (Fig. 3). The differences between the UCE and COI datasets arise due to UCE-based lumping of the following COI-supported morphospecies (Table 1): *S. pezomachiodes* (YMZ056) *+ S. lobatae* (YMZ052), and *S. nr. nigriceps*-2 (YMZ041) + *S. nr. occidentalis* (YMZ021). *Sycophila foliatae* also differs in the UCE data as it was recovered from two separate clades, once grouped with *S. nr. foliatae*-1 as mentioned above (YMZ025), and again as sister to *S. nr. lanae* (YMZ031/32). Nearly all nodes within the UCE dataset are strongly supported by ultrafast bootstraps and SH-aLRT (Fig. S4). Putative species were grouped together regardless of sampling location (e.g. *S*. nr. *dubia/globuli* from CA and IA, *S. texana* from FL and TX), thus ruling out potential phylogeographic substructures at the population level biasing accurate species delimitation. Five of the six species groups from the COI dataset were recovered, with the exception of MOTU10 *Sycophila* sp3 which failed to generate UCE data.

**Figure 3.**
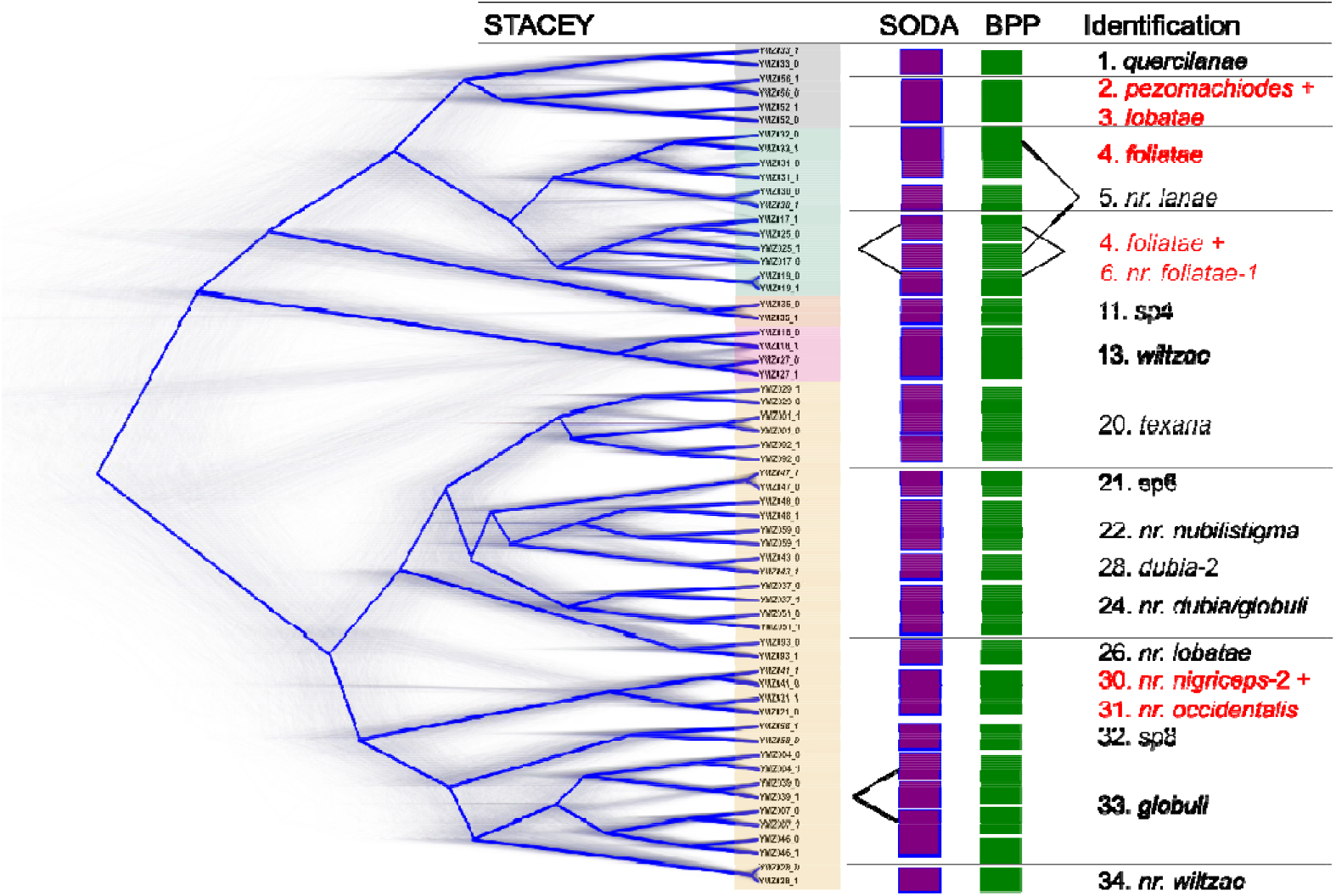
Overview of all UCE data used in inferring *Sycophila* diversity associated with North American oak galls. Left: Allelic phased UCE phylogeny of *Sycophila* using STACEY. SODA and BPP columns indicate assignments of individuals into groups by these respective algorithms. ‘Identification’ describes putative species assignments based on the sum of information to the left of this column. Specimen code “_0” and “_1” represent the phased alleles of the same individual. Topological discordances from the COI data are shown in red. Black lines on the SODA/BPP columns indicate cases where phased alleles did not group as sisters to each other. Colored clades correspond to species groups.

### 3.3 PCoA

We found no correlation between pairwise distances in interaction trait space and pairwise phylogenetic distances between *Sycophila* species pairs were (Mantel *r* = -0.00791, *p* = 0.541, Fig. S5), i.e., both closely-related and distantly related *Sycophila* species (from different species groups) can attack galls with similar defensive trait combinations (Fig. 4A). Sets of unrelated *Sycophila* interact with galls of different size, with most attacking medium and larger galls (Fig. 4B). In terms of external gall defensive traits, *Sycophila* interact more often with galls with minimal external defenses (smooth or textured, Fig. 4C). Additionally, unrelated *Sycophila* commonly attack galls with different internal traits (fleshy, woody, or hollowFig. 4D).

**Figure 4.**
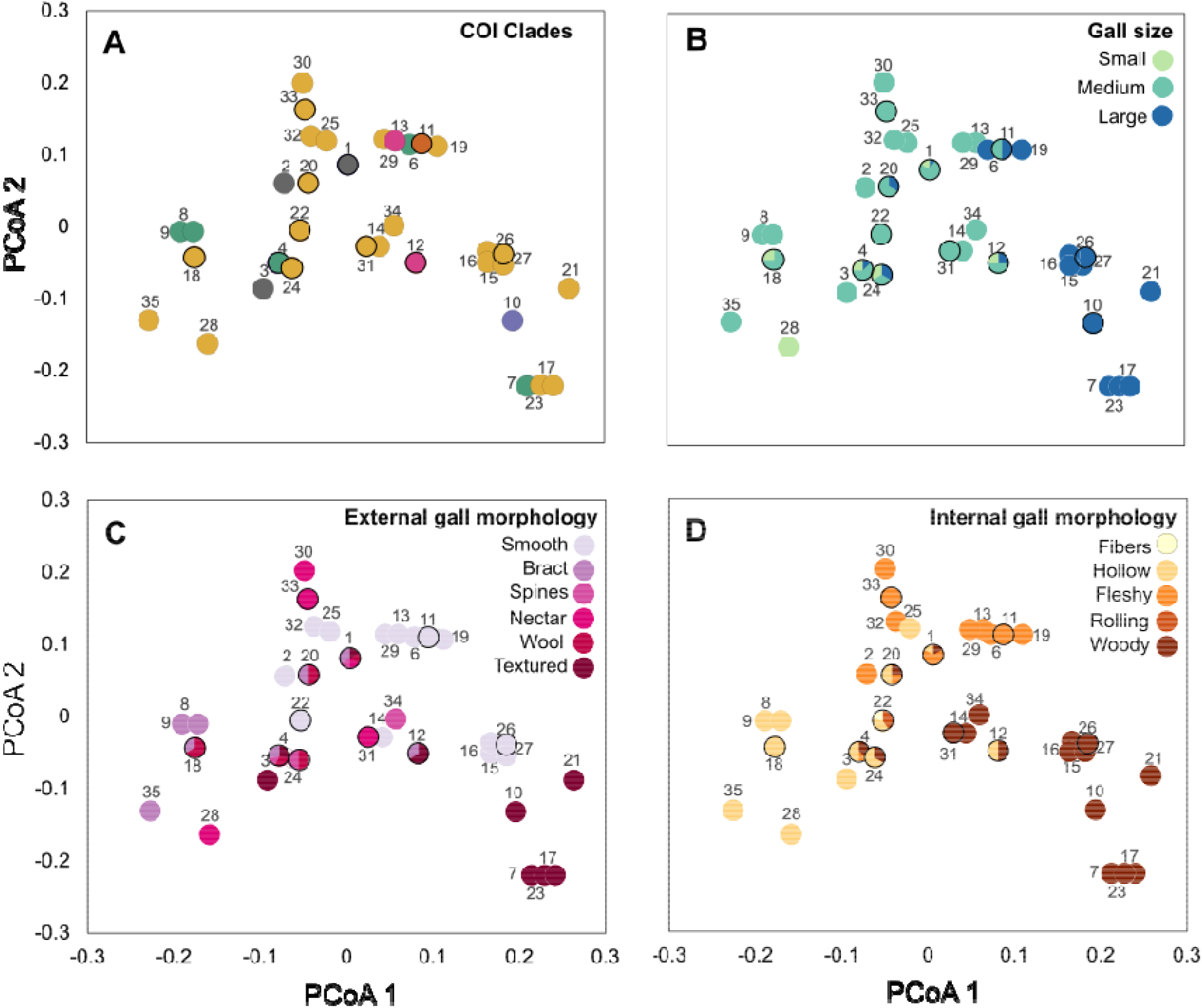
A) Biplots of PCoA1 and PCoA2 showing the centroids of the host gall trait combinations that each *Sycophila* species (numbered) interacts with in gall wasp trait space. Numbers represent *Sycophila* species designated in Table 1. Dark borders around circles represent species reared from more than one host gall. Different colored symbols of *Sycophila* species represent A) COI clades (see Fig. 2 for colors), B) sizes of host galls C) external traits of host galls, D) internal traits of host galls..

## 4. Discussion

Hymenopteran parasitoids are likely one of the most diverse groups of animals (Forbes et al., 2018), yet much of their biology and ecology remains unknown due to their small size and often problematic taxonomy. We used an integrative approach to identify 35 *Sycophila* parasitoids associated with a subset of North American oak gall wasps, a crucial first step for understanding the tritrophic interactions and community assemblage of this species-rich but understudied system. Our work corroborates similar studies within gall systems where generalist species that are thought to have wide host breadth and geographic ranges have been revealed to be a suite of cryptic specialists (e.g. Kaartinen et al, 2010; Sheikh et al, 2022). This of course has direct consequences such as understanding the effectiveness of these parasitoids as potential biocontrol agents (Davis et al., 2018; Smith-Freedman et al., 2019). With this refined dataset on the host ranges and preference of each species, we can more accurately identify the host traits that define, or are components of, parasitoid niches, and thus gain insights into axes that are relevant for structuring tritrophic interactions.

### 4.1 *Sycophila* host specificity

We used a combination of molecular data (COI, UCE) and extensive ecological data to determine the species richness and host repertoire of *Sycophila* parasitoids associated with oak cynipid galls in North America. Based on our conservative delimitations of potential species, most *Sycophila* are oligophagous species that have limited host repertoire across host tree relationships and gall morphology (Table 1, Figs. 2, 3). For example, 11/17 species represented in the UCE phylogeny were only found on one oak Section (Fig. 3), while certain species (*S. nr. nigriceps/nr. occidentalis/*sp4/*globuli*) were reared from galls from multiple oak Sections but with similar morphology (e.g., all woody stem galls). The true host breadths of some or all of these *Sycophila* species are likely higher given that we did not sample from all of the 700+ described North American oak gall wasp species, nor did we sample across the entire range of the included species, and we still don’t know the full cynipid diversity in North America. Unlike the host gall wasps, which are largely restricted to inducing galls on related oak trees and often in the same oak Section (Cook et al., 2002; Stone et al. 2009; Melika et al. 2010; Tang et al., 2011), at least some *Sycophila* species specialize on aspects of the gall itself. This result is consistent with previous works examining host traits of another koinobiont endoparasitoid *Euderus set* (Eulophidae), which is reared from distantly related gall wasp genera from different oak tree Sections, but apparently only successfully attacks integral leaf and stem galls lacking external defenses (Ward et al., 2019).

In terms of what these data reveal about *Sycophila* overcoming host gall defenses, most *Sycophila* species (20/35) were reared from galls with minimal external defenses and a variety of different internal gall textures (Figs. 2, 4C, 4D). By comparison, galls with external spiny, wooly, or nectar-secretions were attacked only by a smaller subset (13/35) of *Sycophila* species (Fig. 4C). Similarly, galls with internal defenses such as radiating internal fibers or with free-rolling larval chambers, as seen in cynipid galls of some *Amphibolips* and *Dryocosmus* spp. in our study, were only attacked by a single species of *Sycophila*, whereas the woody or fleshy internal morphologies in typical galls were attacked by multiple species (Fig. 4D). This suggests that some external or internal gall traits serve to reduce attack by, or wholly exclude, some *Sycophila* parasitoids. The PCoA biplot and results of the mantel test also showed that associations between *Sycophila* and specific gall trait combinations might have evolved convergently as distantly related species are found attacking galls with similar defenses. However, without factoring in the pattern of extended phenotype evolution (i.e. convergence of gall structure), we cannot distinguish whether the underlying process is one of codiversification/cospeciation, or one of host switching. Unfortunately, despite recent advances in understanding of basal relationships between tribes within the Cynipidae (Blaimer et al., 2020; Brandão-Dias et al., 2022; Ward et al., 2022; Zhang Y.M. et al., 2020) and work on relationships between gall wasp lineages in the Western Palearctic (Stone et al. 2009), the status of oak gall wasp taxonomy and phylogenetics in North America is incomplete and many genera are paraor even polyphyletic. Hopefully with the ongoing research of global and North American Cynipini phylogeny this caveat can be addressed in the near future.

Additionally, the general patterns of *Sycophila* host preference listed above do not account for interactions with other natural enemies within the gall system including hyperparasitoids, which can target and kill mature *Sycophila* larvae, and therefore affect the patterns we observe in terms of adult emergence. Unfortunately, many of the species interactions within North American oak gall communities remains unknown, aside from *Ormyrus* (Sheikh et al., 2022), but we hope these studies will lay the foundations for, and generate interest in, future investigations that can clarify these complex community structures. Future studies could be conducted focusing on intensive sampling at a smaller geographic scale, to help clarify whether the patterns we observed in our dataset are influenced by sampling bias. As the galls were often collected haphazardly based on availability, the rate of parasitism by *Sycophila* (and other parasitoids) cannot be accurately estimated for each of the gall types.

### 4.2 Host Phenology

Another important determinant of a parasitoid’s ability to attack a host is phenology, including the developmental timing of the gall and (or) the seasonal timing of the host plant. For example in our study and other natural enemies within the oak gall systems is the importance of allochronic differentiation, where different species of parasitoid wasps utilize the same host, or a few closely related hosts at different times of the year (Nicholls et al, 2018; Sheikh et al, 2022; Zhang L. et al., 2019). The optimal temporal window for oviposition into a particular species of gall may often be limited to the time before the gall grows too large for ovipositors to reach the insect inside. Based on the PCoA analysis (Fig. 4B) more *Sycophila* species attacked medium and larger galls, which is surprising given the relatively shorter ovipositor length when compared with other parasitoids such as *Torymus*. This suggests the oviposition by *Sycophila* must occur when the galls are early in the developmental stage, which is often a narrow window of time during the oak leaf flushing that are often species-specific (Zhang L. et al. 2019). The alternative explanation being that some *Sycophila* species are targeting inquilines instead of gall inducers, but without detailed dissection studies this cannot be verified. The majority of our *Sycophila* were collected from asexual generation galls found in autumn (27/44), which are often more conspicuous and have a longer growing period, compared to sexual generation galls (17/44) which often develop rapidly in spring on ephemeral resources such as catkins. Nevertheless, we did rear some *Sycophila* species only from the sexual generations (*S*. nr. *lanae, S*. nr. *foliatae*-1, *S. wiltzae, S*. sp5-1, *S*. sp5-2, *S. flava*, and *S. texana*) or from galls from both generations (*S. foliatae*, nr. *lobatae*-2, *S*. sp3, *S*. sp4, *S. marylandica*, and *S*. nr. *dubia/globuli*). Some of these parasitoids might therefore be bi-or multivoltine, having multiple generations a year attacking different galls at various stages of development (Askew 1965). Bivoltinism is known for several of the chalcids attacking European oak cynipid galls, including species in which the two generations have different ovipositor lengths, allowing them to attack different gall morphologies (Askew 1965). Studies have also shown that the emergence phenology of sympatric gall wasp populations can differ based on phenological differences between host plants, which can reduce gene flow between host-associated populations (Hood et al., 2019; Zhang L. et al., 2019). While some studies have shown that temporal isolation can cascade across multiple trophic levels and potentially drive the speciation of some parasitoid communities (Hood et al., 2015, Zhang L. et al. 2019), the study by Sinclair et al. (2015) showed that different oak galls respond differently to variation in host phenology, and that being a generalist requires maintaining phenological flexibility.

### 4.3 Species delimitation and taxonomy of *Sycophila*

The resolution offered by UCE data is promising for generating robust phylogenies at the species/population level, especially with allelic phasing and the extraction of SNPs (Andermann et al., 2019; Gueuning et al 2020; Prebus 2021). Effects of potential gene flow, incomplete lineage sorting and/or introgression can be seen in the form of incongruencies within the UCE trees (Fig. 2, *S. globuli, S. nr. foliatae*-1, *S. nr. lanae, S. foliatae*), as the two alleles of the same sample were not recovered as sisters to each other. We acknowledge the potential inflation of putative species richness based on the molecular species delimitation methods used (Chambers & Hillis, 2020; Luo et al., 2018; Sukumaran & Knowles, 2017), especially when there are no clear barcoding gaps in some species (eg. 4.3% intraspecific divergence within *S. flava/texana*, while *S*. nr. *foliatae*-2 and *S*. nr. *lanae* have only 3.9% interspecific divergence). UCE loci have been shown to be useful for species delimitation in some Hymenoptera (Branstetter & Longino, 2019; Gueuning et al., 2020; Longino & Branstetter, 2021), and in our study to be more conservative than the traditional DNA barcodes as multiple COI morphospecies were lumped together based on UCE results (Figs. 2, 3). However, our exploration using phased SNPs and a subset of UCE loci using various delimitation software corroborates findings from other phylogenomic species delimitation studies that some taxa can remain contentious (Prebus, 2021; Samacá-Sáenz et al., 2020). It is likely that the discordance within our UCE data such as the *S. nr. foliatae*-1*/S. nr. lanae/S. foliatae* clade is the result of over-splitting and might represent a single variable species, introgression, or a recent/ongoing divergence. Future studies should focus on wider geographic sampling for these challenging complexes using a population genomic approach to detect geographical substructures and/or ongoing gene flow (Bunnefeld et al., 2018).

Nevertheless, it is clear that the North American *Sycophila* is in need of taxonomic revision, and while this is beyond the scope of the current study, the molecular evidence presented here and in previous studies (Davis et al., 2018; Smith-Freedman et al., 2019) has shown that body coloration or wing band shape, as used by Balduf (1932), can vary significantly among conspecifics, and are therefore not reliable diagnostic characters. This is especially evident in species with a small wing band (e.g. *S. quercilanae, S. pezomachiodes, S. marylandica, S. wiltzae*), where the females have seemingly diagnostic color patterns, but the males look nearly identical and cannot be identified. Thus, a thorough exploration of morphological, ecological, and biogeographic data combined with phylogenomic data and more complex species delimitation methods are needed to be able to determine the species limits within the genus *Sycophila*. Additional studies on the biology of different North American *Sycophila* species could potentially explain the difference between host repertoires, as this genus includes both endoparasitoids (Claridge 1959; Gómez et al., 2013), which are often specialists due to the need to overcome host immune defenses, and ectoparasitoids (Gates et al., 2020), that are more often generalists.

## Acknowledgements

We would also like to thank Maureen Turcatel, Bernardo Santos, Karen Neves, and Matt Prebus for providing time and expertise for UCE library preparation and downstream analyses. We acknowledge University of Florida Research Computing (http://researchcomputing.ufl.edu/) and the Smithsonian Institution High Performance Cluster (https://doi.org/10.25572/SIHPC) for providing computational resources and support that have contributed to the research results reported in this publication. YMZ was funded by the Theodore Roosevelt Memorial Grant provided by the American Museum of Natural History and Oak Ridge Institute for Science and Education (ORISE) fellowship. Funding to AKGW was awarded by the American Genetic Association. GNS is funded by the UK NERC Discovery grant NE/T000120/1. Mention of trade names or commercial products in this publication is solely for the purpose of providing specific information and does not imply recommendation or endorsement by the USDA. USDA is an equal opportunity provider and employer.

## Author Contributions

Y.M.Z., S.I.S., A.K.G.W., and A.A.F. designed the study. All authors made collections and/or reared animals. Y.M.Z., S.I.S., A.K.G.W., and C.D. obtained the sequence data. Y.M.Z., S.I.S., A.K.G.W., A.A.F., and K.M.P. conducted the analyses, Y.M.Z., S.I.S., A.A.F., K.M.P., and G.N.S. wrote the manuscript. All authors contributed to specimens, revisions and approved the final manuscript.

## Data Availability Statement

The COI data is available on GenBank (MZ905524–905639), raw UCE sequences available on SRA (SAMN20307313–20307342), for full details see Table S1. COI and UCE alignment files, and input file for STACEY/BPP are available on the Dryad Digital Repository at: https://doi.org/10.5061/dryad.x0k6djhmb.

---

**Figure S1**. Summary and identification of *Sycophila* included in this study.

**Figure S2**. Photos of galls included in this study. All photos by authors except for *Callirhytis perdens* by Leslie Flint, *Disholcaspis edura* by Mike Plagens, and *Zapatella davisae* by Kelly Omand.

**Figure S3**. Full COI phylogeny of *Sycophila* associated with oak galls. Nodal support represents % of bootstrap pseudoreplicates.

**Figure S4**. Full UCE phylogeny of *Sycophila* associated with oak galls. Dots at the nodes represent strong support for Ultrafast Bootstrap (≥ 95) and SH-aLRT (≥80).

**Figure S5**. Top. Biplots of PCoA1 and PCoA2 for host galls plotted in gall wasp trait space. Bottom. Scatter plot of net evolutionary divergence and distance in interaction trait space between each pairwise combination of *Sycophila* species.

**Table S1**. Collection information for *Sycophila* samples, including GenBank/SRA accession numbers.

**Table S2**. A table of the gall morphological traits.

**Table S3**. Intraspecific divergence of *Sycophila* COI species.

**Table S4**. Interspecific divergence of *Sycophila* COI species.

**Table S5**. Gall wasp and *Sycophila* interaction matrix.

