## Supplementary material for "Delimiting the cryptic diversity and host preferences of *Sycophila* parasitoid wasps associated with oak galls using phylogenomic data": Fig. S1

**SUPPLEMENTAL FIGURE 1. *Sycophila* morphospecies**

**Species 1. *Sycophila quercilanae***

**Host galls: *Acraspis erinacei, Ac. macrocarpae, Andricus quercusfrondosus, An. quercuspetiolicola, Callirhytis favosa, Disholcaspis quercusglobulus, D. quercusmamma, Kokkocynips imbricariae, Melikaiella ostensackeni, Me. tumifica, Neuroterus quercusbatatus,* and *Philonix nigra*.**

**Host plants: *Q. alba, Q. bicolor, Q. falcata, Q. macrocarpa, Q. palustris, Q. rubra*, and *Q. velutina*.**

**Locality: IA, KY, MO, and TN**

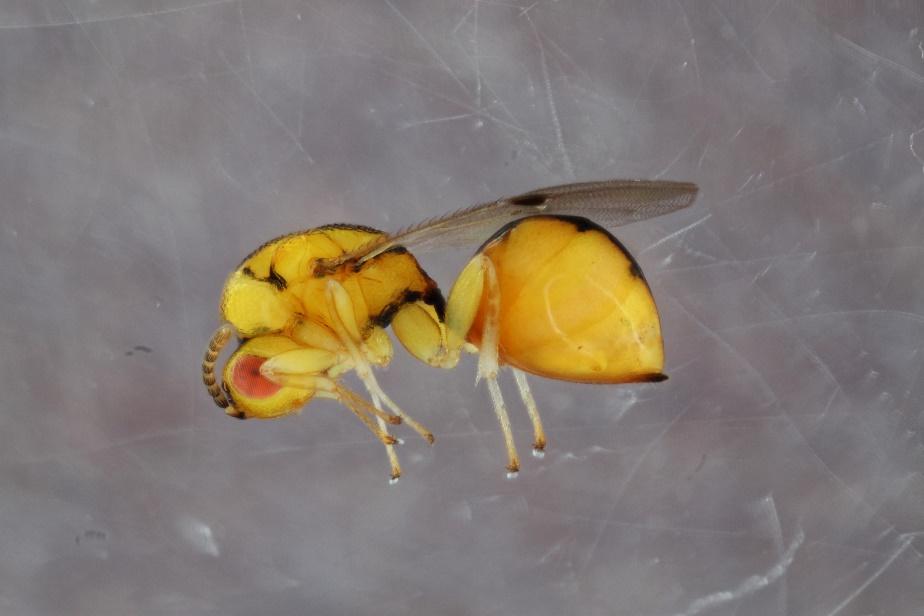

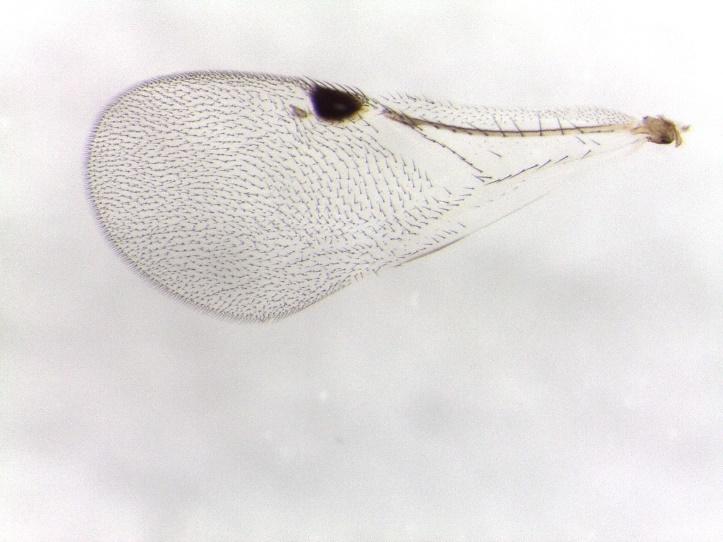

**Species 2. *Sycophila pezomachiodes***

**Host gall: *Disholcaspis edura***

**Host plant: *Quercus arizonica***

**Locality: AZ**

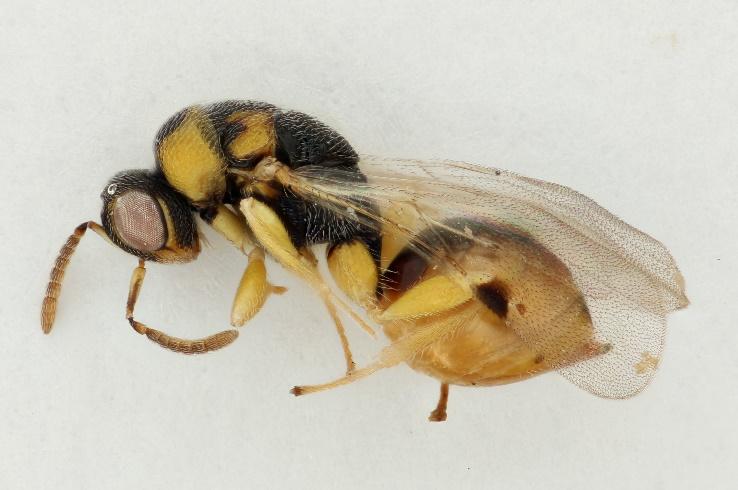

**Species 3. *Sycophila lobatae***

**Host gall: *Heteroecus sanctaeclarae***

**Host plant: *Quercus chrysolepis***

**Locality: CA**

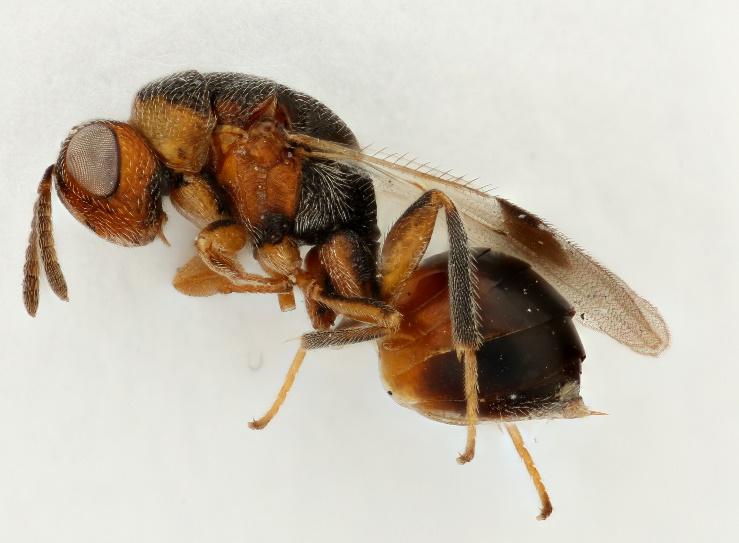

**Species 4. *Sycophila foliatae***

**Host galls: *Acraspis erinacei, Ac. macrocarpae, Ac. pezomachiodes, Andricus ignotus, An. quercusflocci, An. quercusfoliatus, An. quercusfrondosus,* and *Bassettia pallida*.**

**Host plant: *Quercus alba, Q. geminata, Q. macrocarpa,* and *Q. virginiana.***

**Locality: IA, FL, and MO**

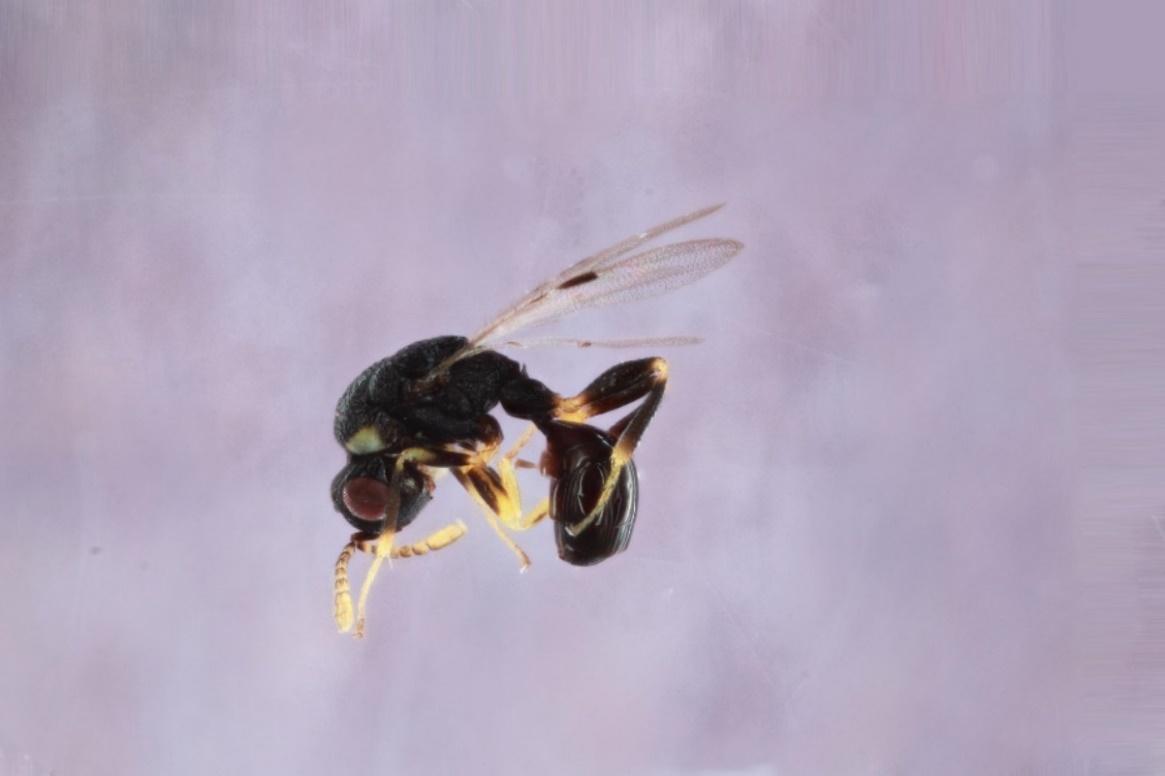

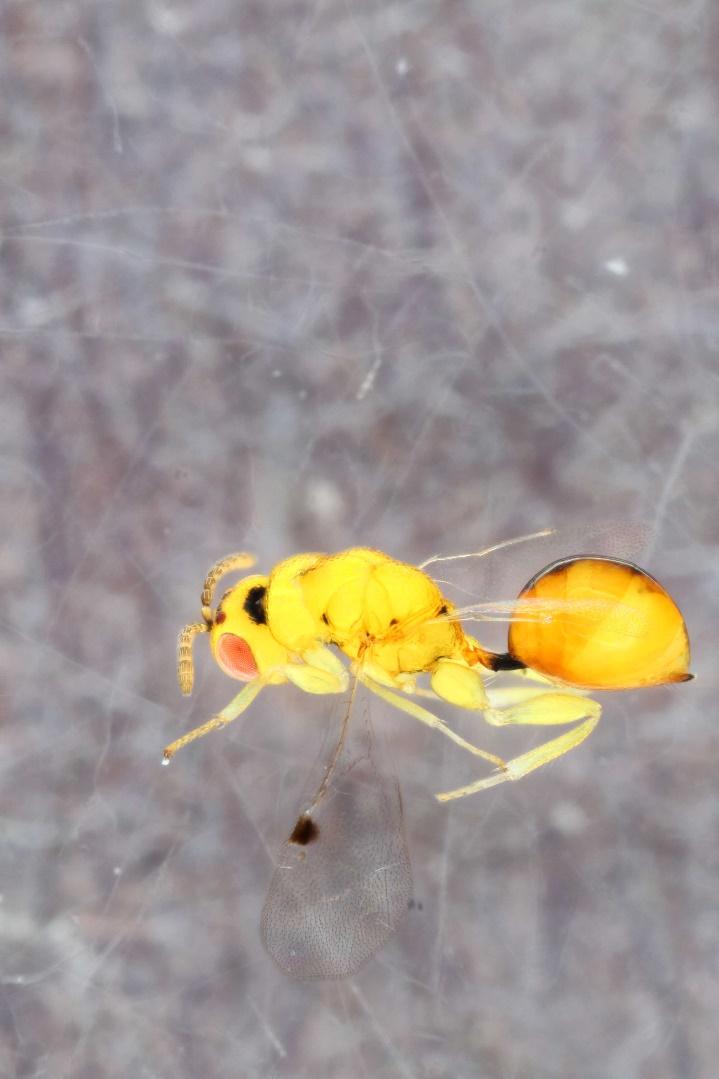

**Species 5. *Sycophila* nr. *lanae***

**Host gall: unidentified acorn gall**

**Host plant: *Quercus rubra***

**Locality: IA**

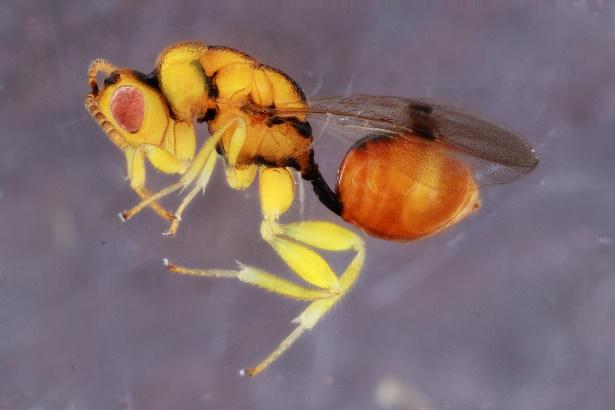

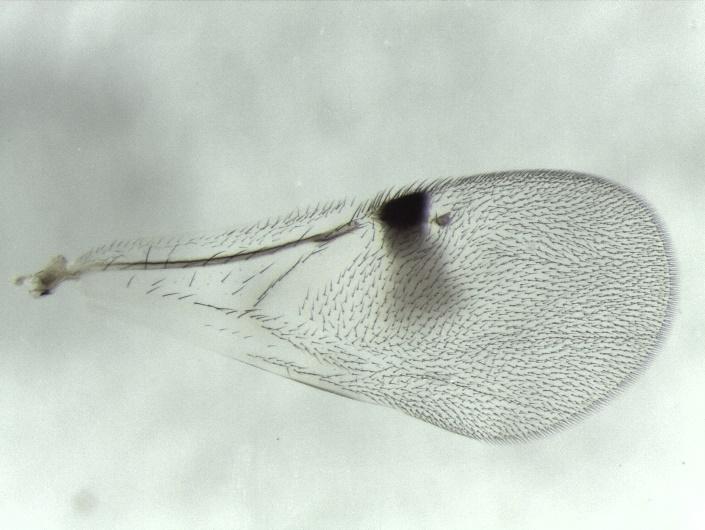

**Species 6. *Sycophila* nr. *foliatae*-1**

**Host gall: *Callirhytis flavipes***

**Host plant: *Quercus bicolor* and *Q. macrocarpa***

**Locality: IA**

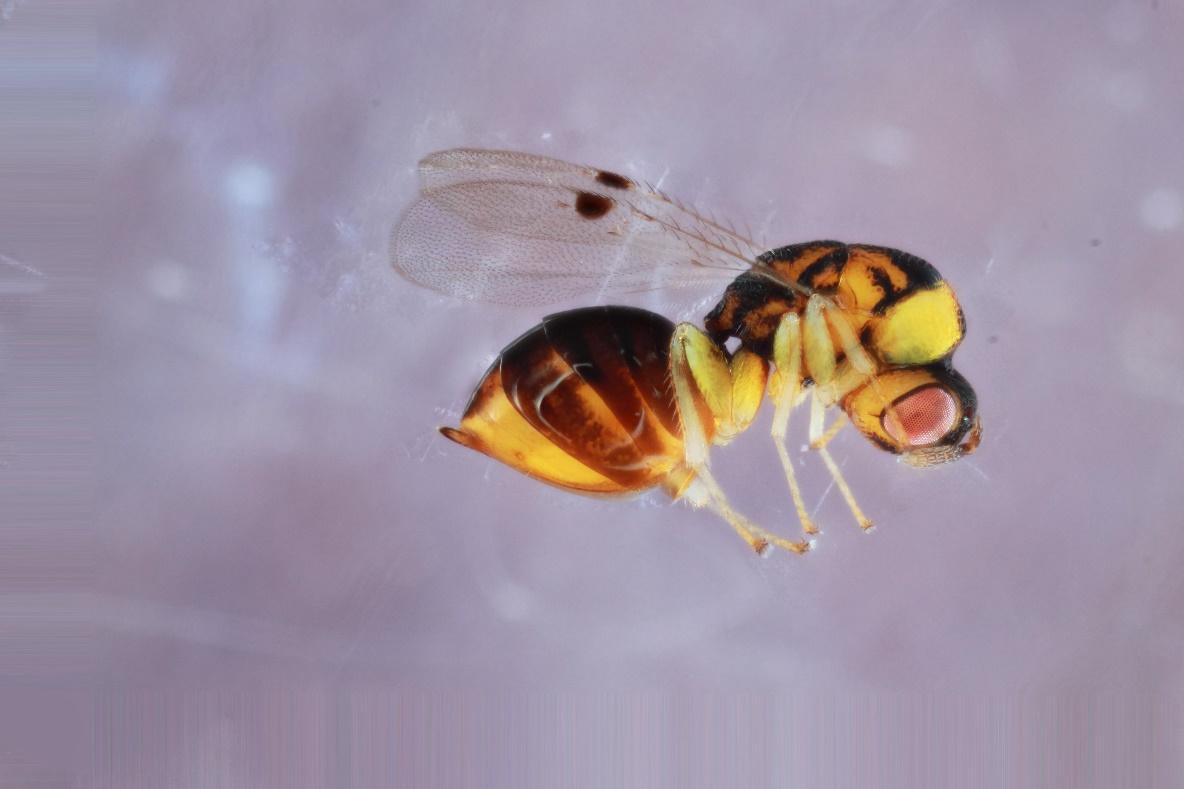

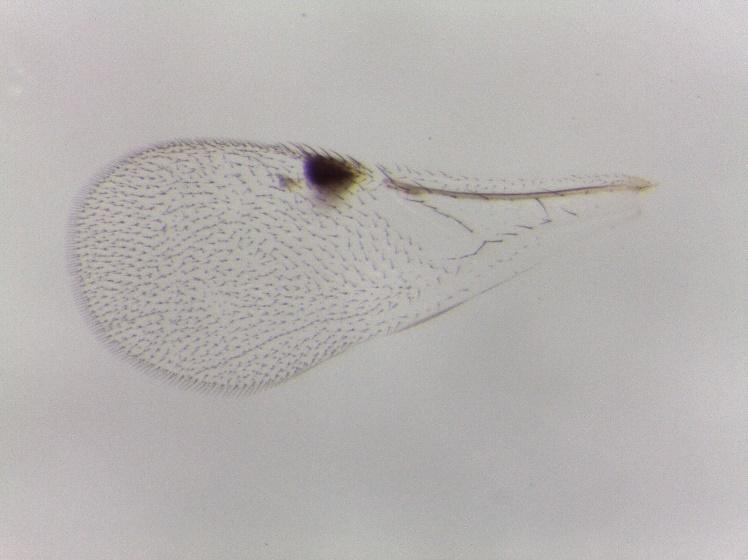

**Species 7. *Sycophila* nr. *foliatae*-2 (see Smith-Freedman et al. 2019 Supplemental Data for images)**

**Host gall: *Zapatella davisae***

**Host plant: *Quercus velutina***

**Locality: MA**

**Species 8. *Sycophila* sp1**

**Host gall: *Andricus quercusfoliatus***

**Host plant: *Quercus geminata* and *Q. virginiana***

**Locality: FL**

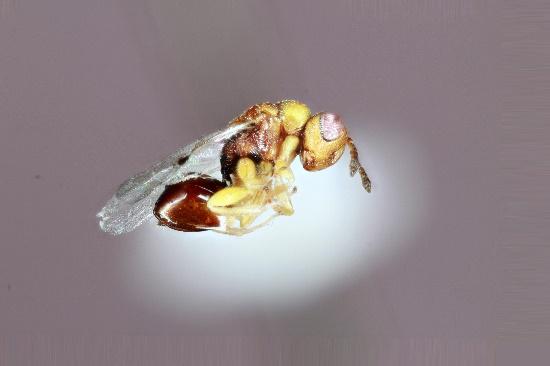

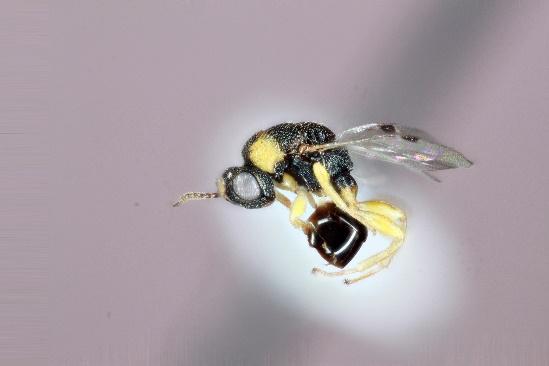

**Species 9. *Sycophila* sp2**

**Host gall: *Andricus quercusfoliatus***

**Host plant: *Quercus geminata* and *Q. virginiana***

**Locality: FL**

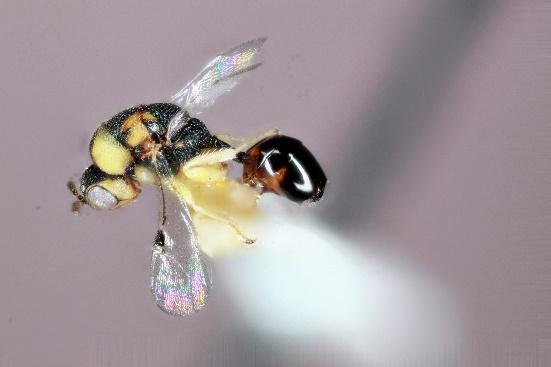

**Species 10. *Sycophila* sp3**

**Host gall: *Callirhytis scitula* and *Zapatella davisae***

**Host plant: *Quercus imbricaria* and *Q. velutina***

**Locality: IA and MA.**

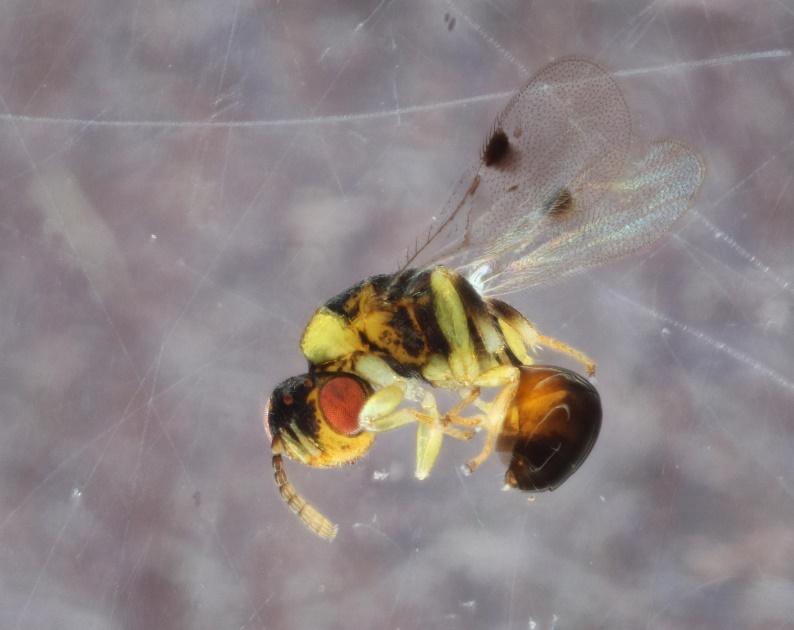

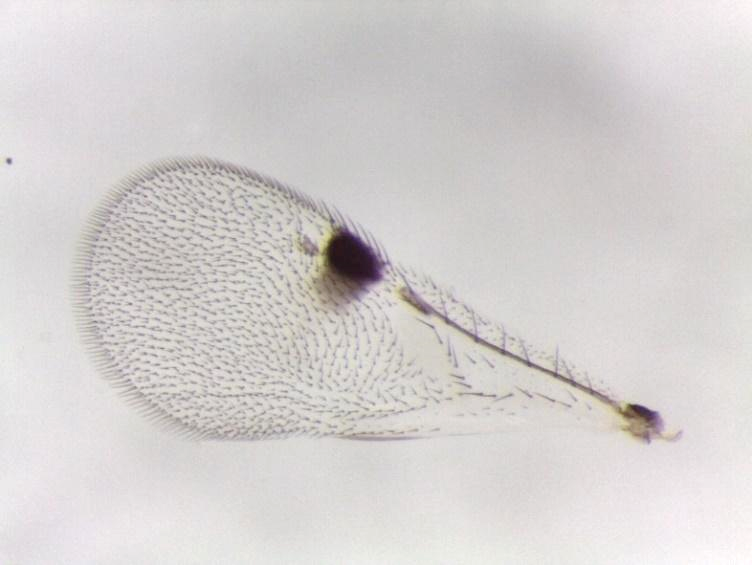

**Species 11. *Sycophila* sp4**

**Host gall: *Melikaiella ostensackeni* and *Neuroterus quercusirregularis***

**Host plant: *Quercus palustris* and *Q. stellata***

**Locality: IA and TX**

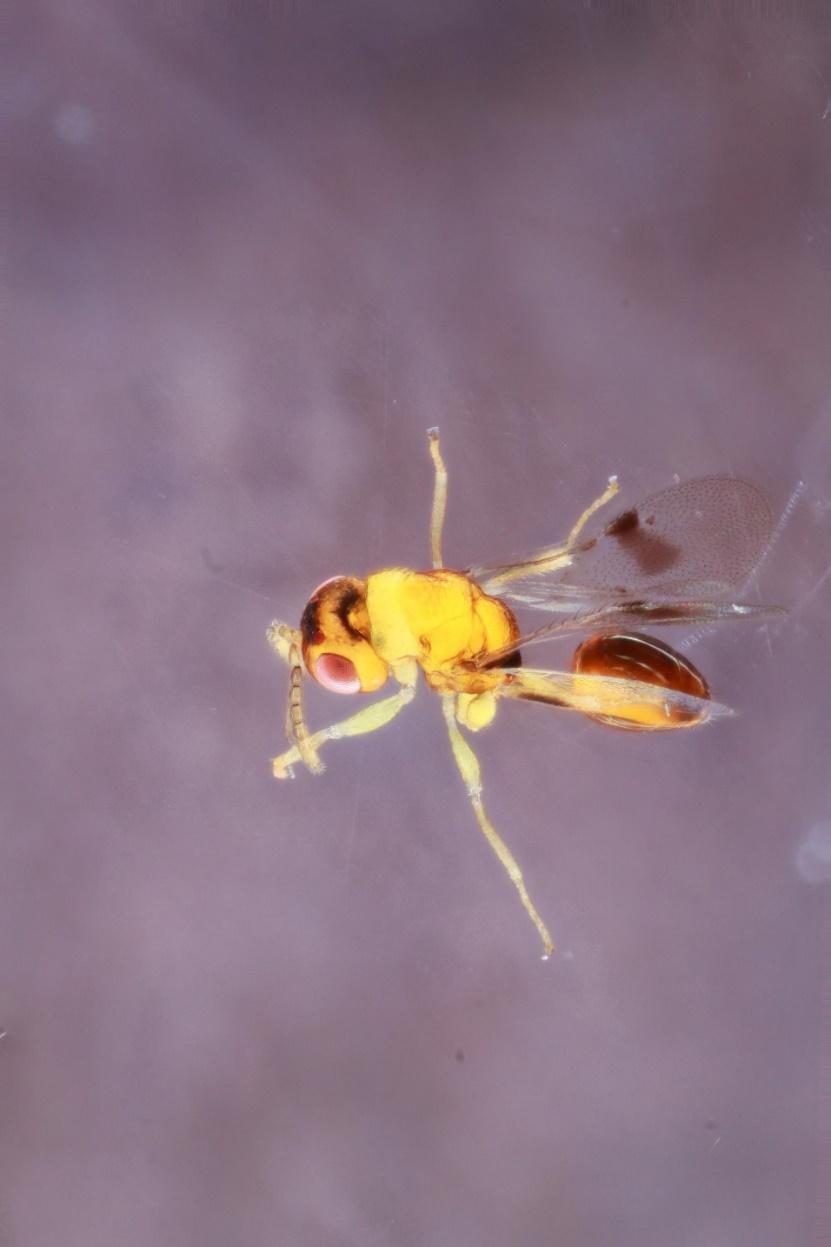

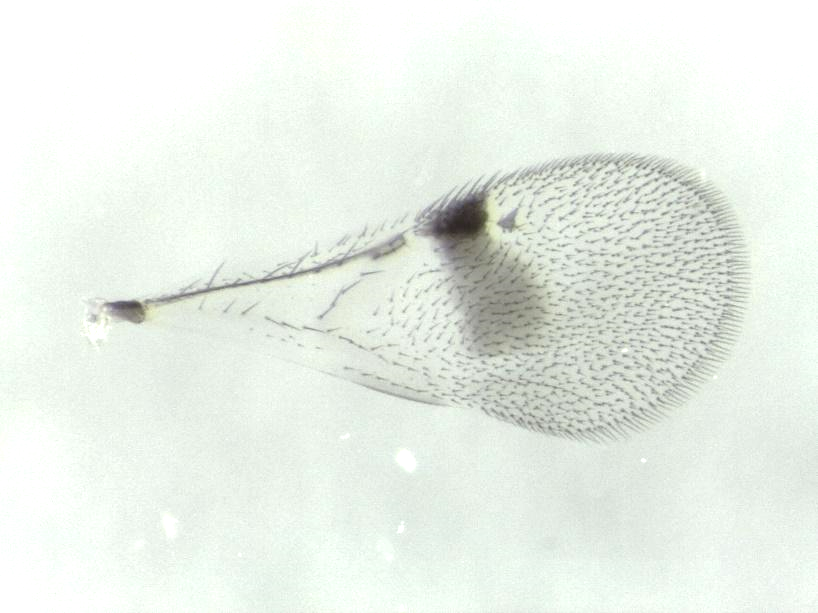

**Species 12. *Sycophila marylandica***

**Host gall: *Acraspis erinacei, Andricus quercuspetiolicola, An. utriculus, Loxaulus quercusmammula*, and *Zapatella davisae***

**Host plant: *Quercus alba, Q. bicolor, Q. palustris*, and *Q. velutina***

**Locality: IA and MA**

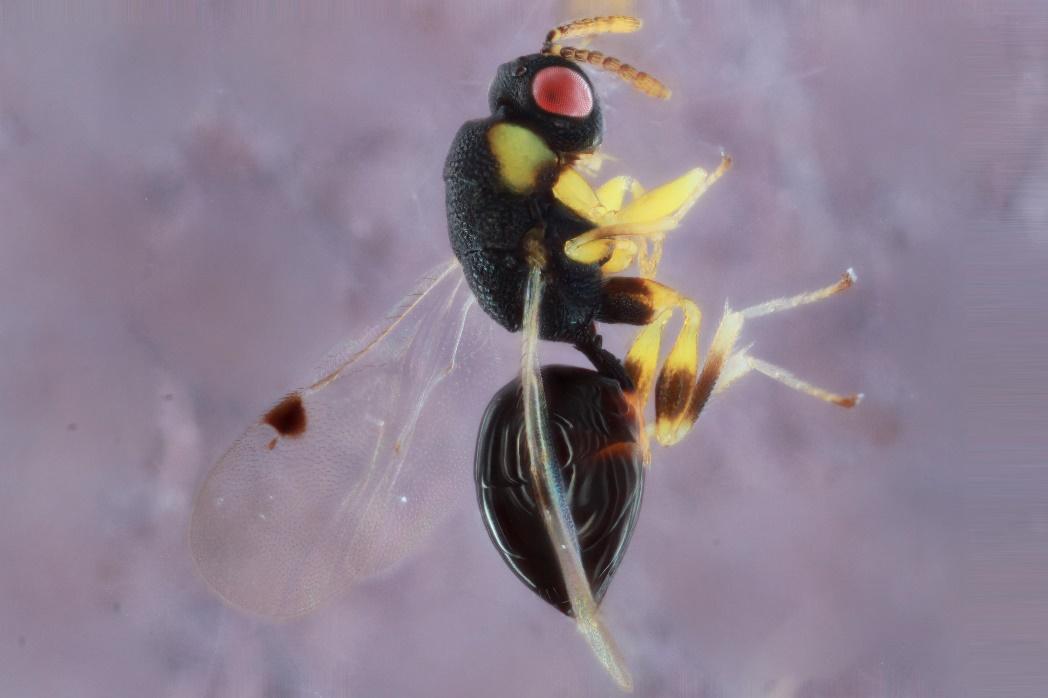

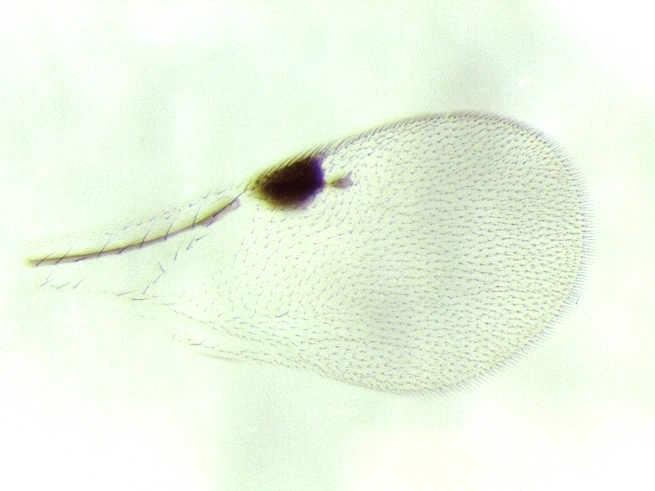

**Species 13. *Sycophila wiltzae***

**Host gall: *Neuroterus washingtonensis***

**Host plant: *Quercus garryana***

**Locality: BC and WA**

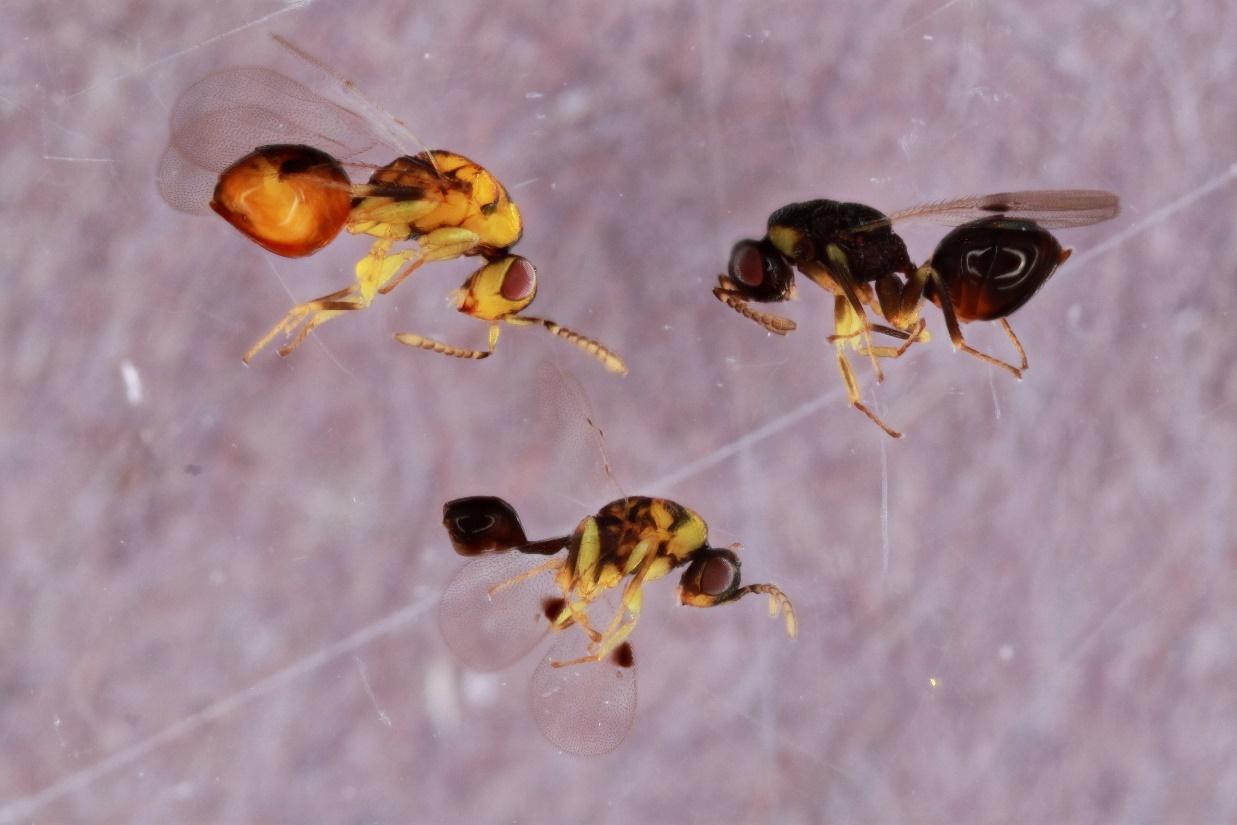

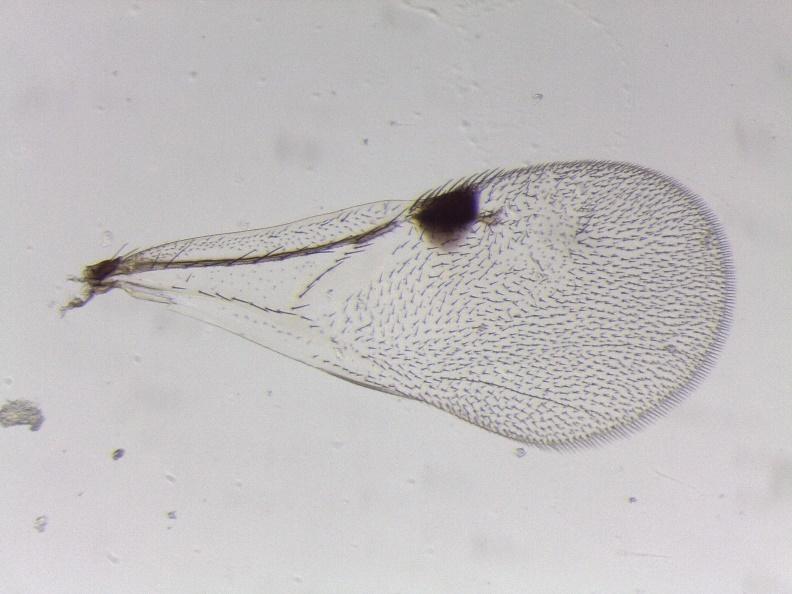

**Species 14. *Sycophila* *varians***

**Host gall: *Kokkocynips imbricariae***

**Host plant: *Quercus falcata***

**Locality: KY**

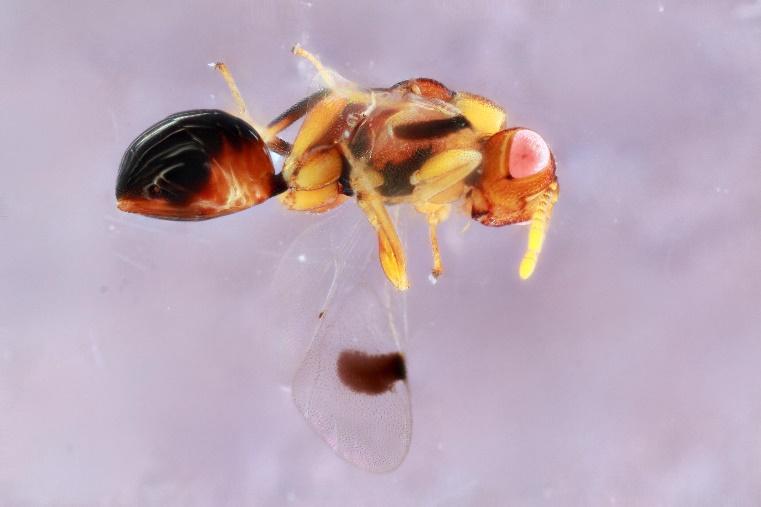

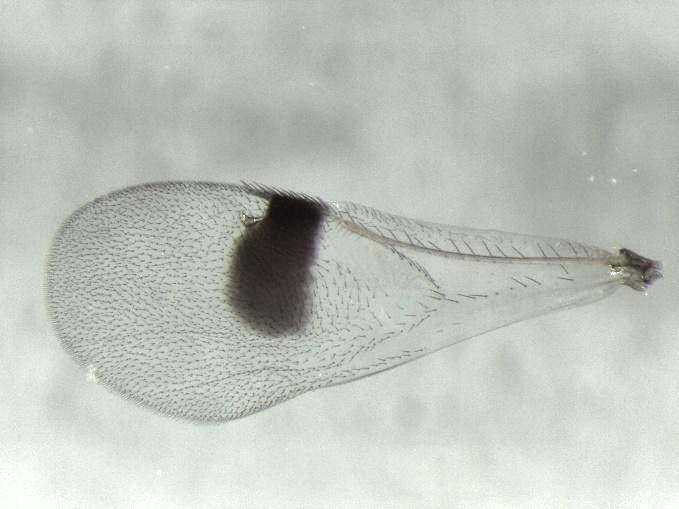

**Species 15. *Sycophila* sp5-1**

**Host gall: *Callirhytis scitula***

**Host plant: *Quercus imbricaria***

**Locality: IA**

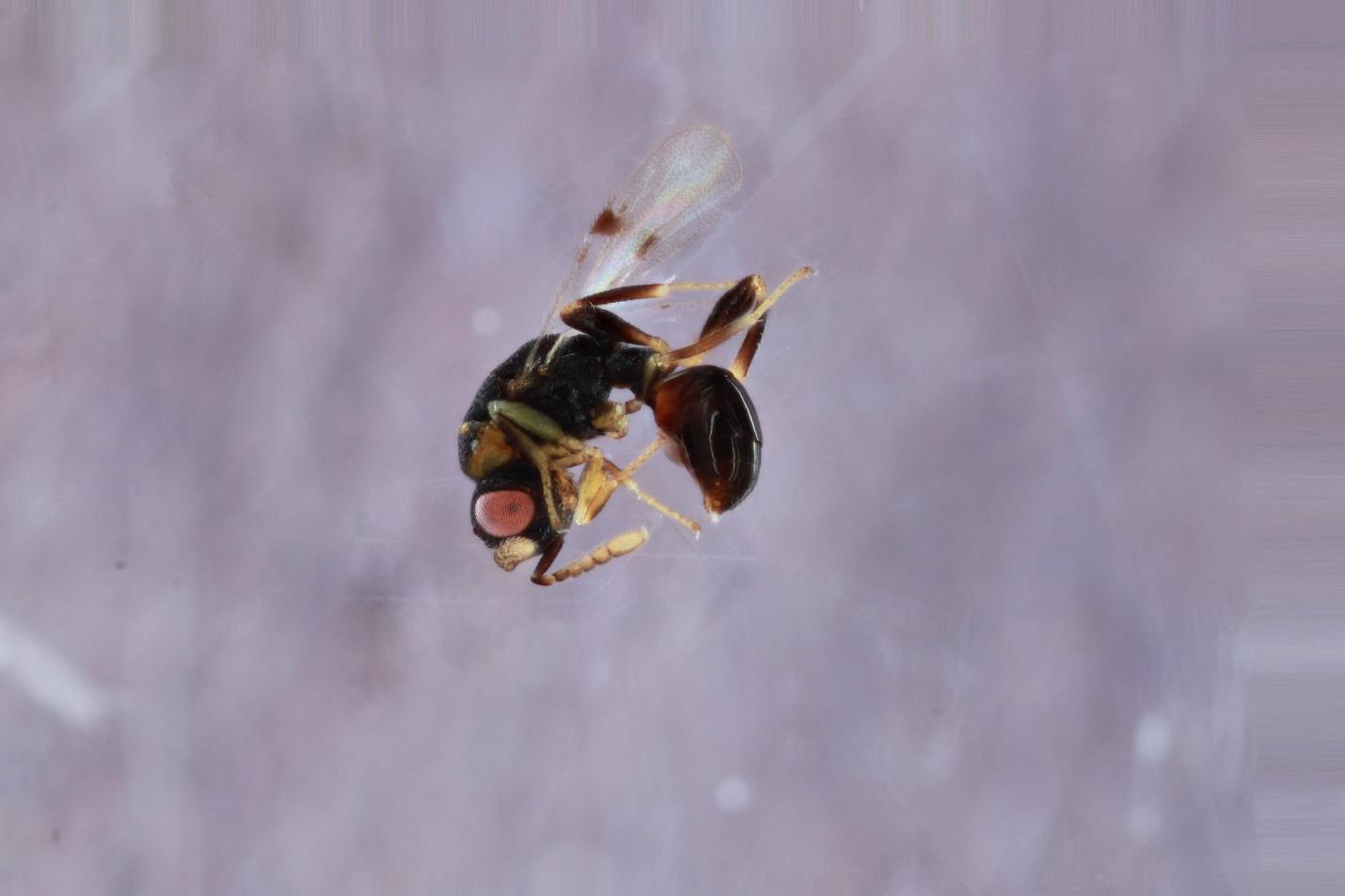

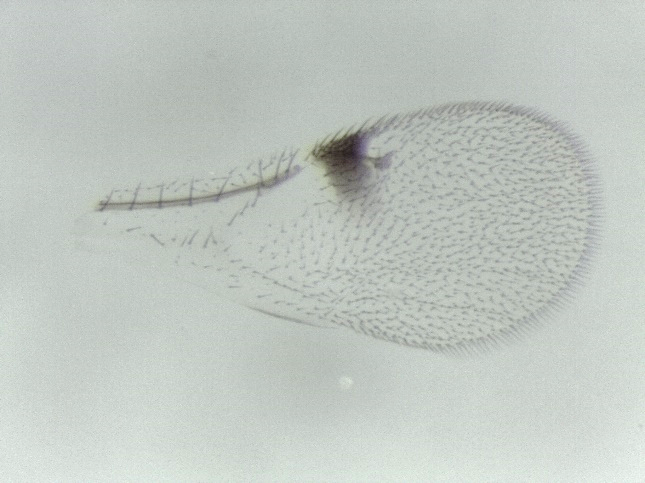

**Species 16. *Sycophila* sp5-2**

**Host gall: *Callirhytis scitula***

**Host plant: *Quercus palustris***

**Locality: IA**

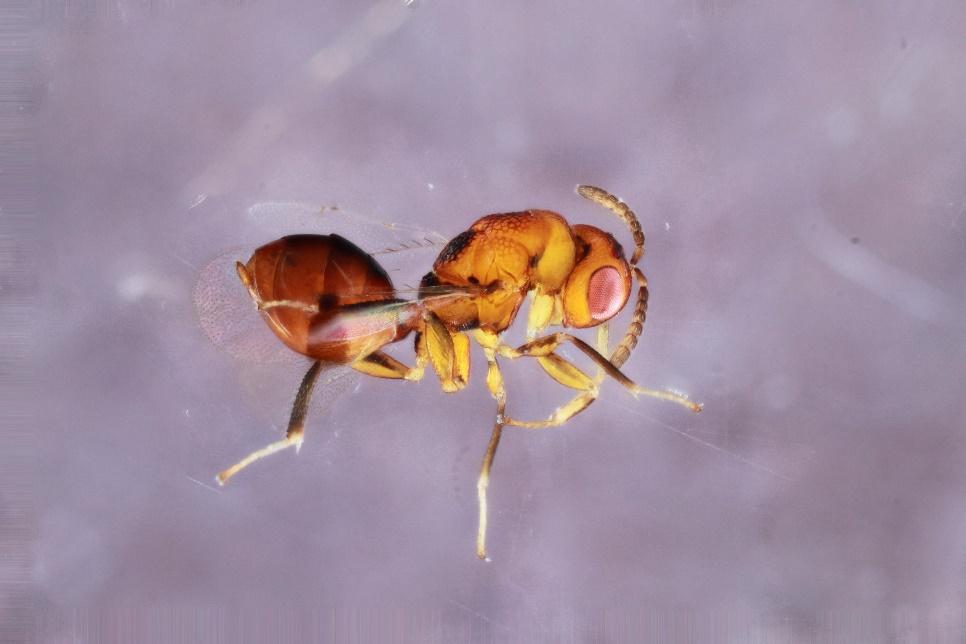

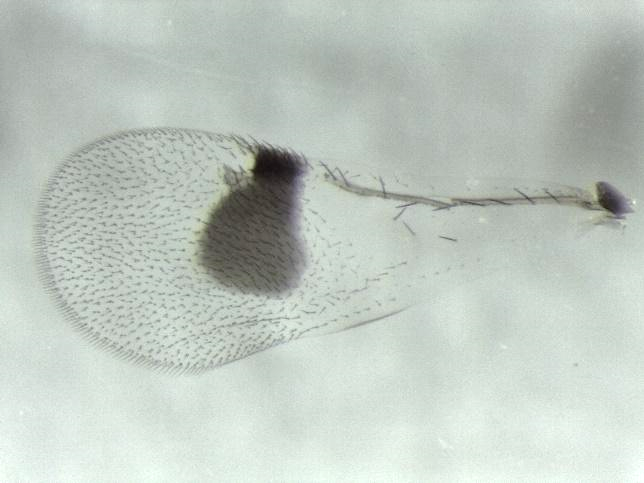

**Species 17. *Sycophila nr. flava* (see Smith-Freedman et al. 2019 Supplemental Data for images)**

**Host gall: *Zapatella davisae***

**Host plant: *Quercus velutina***

**Locality: MA**

**Species 18. *Sycophila flava/texana***

**Host gall: *Andricus ignotus, An. quercusfoliatus, An. quercuslanigera*, and *Belonocnema kinseyi***

**Host plant: *Quercus geminata, Q. macrocarpa*, and *Q. virginiana***

**Locality: IA, FL, and TX**

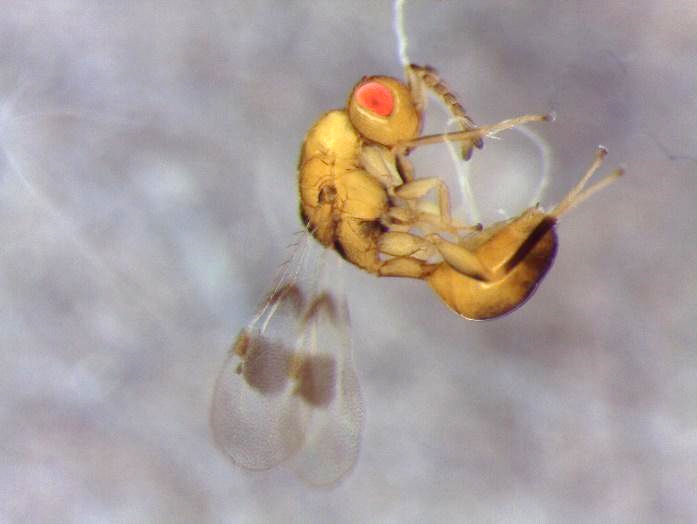

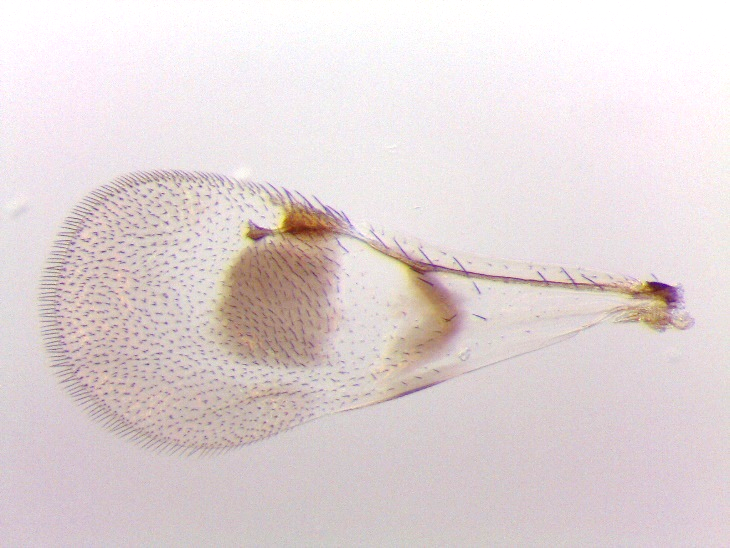

**Species 19. *Sycophila flava***

**Host gall *Neuroterus quercusbatatus***

**Host plant: *Quercus bicolor***

**Locality: IA**

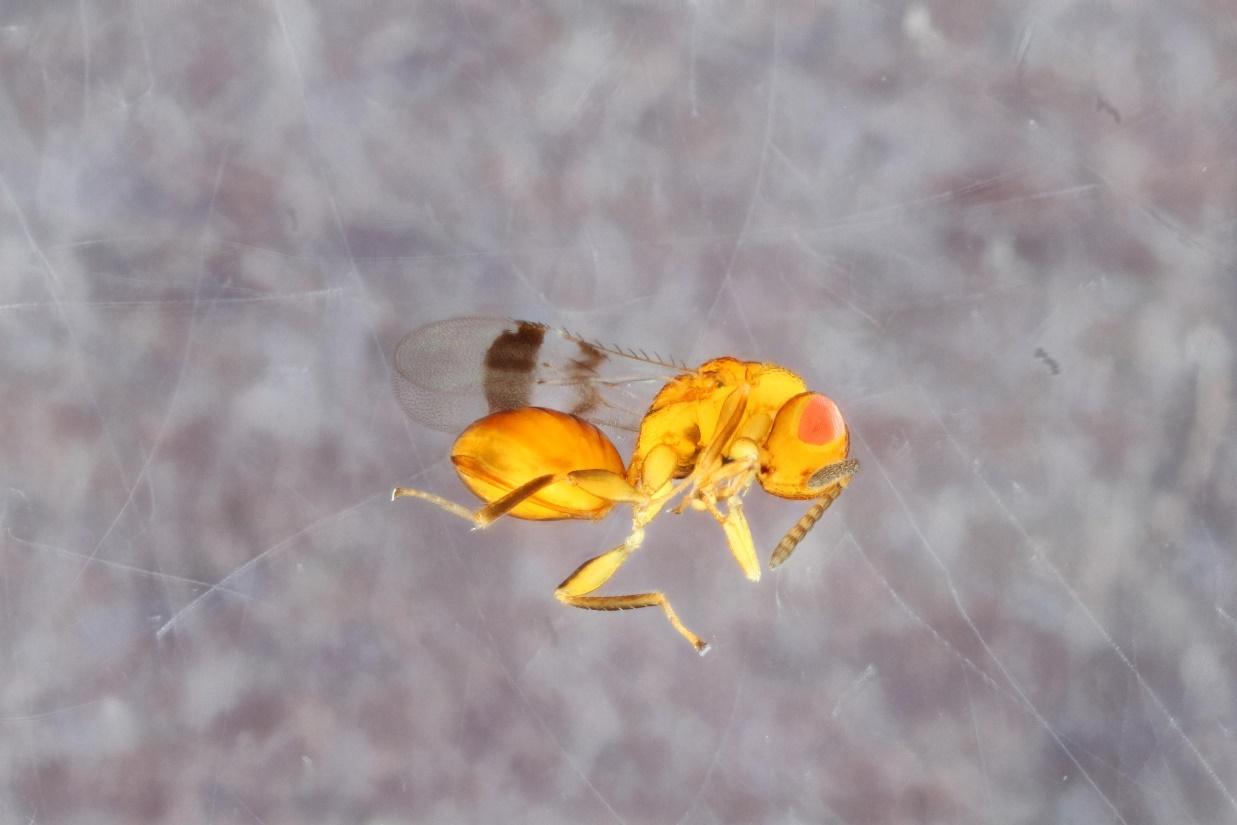

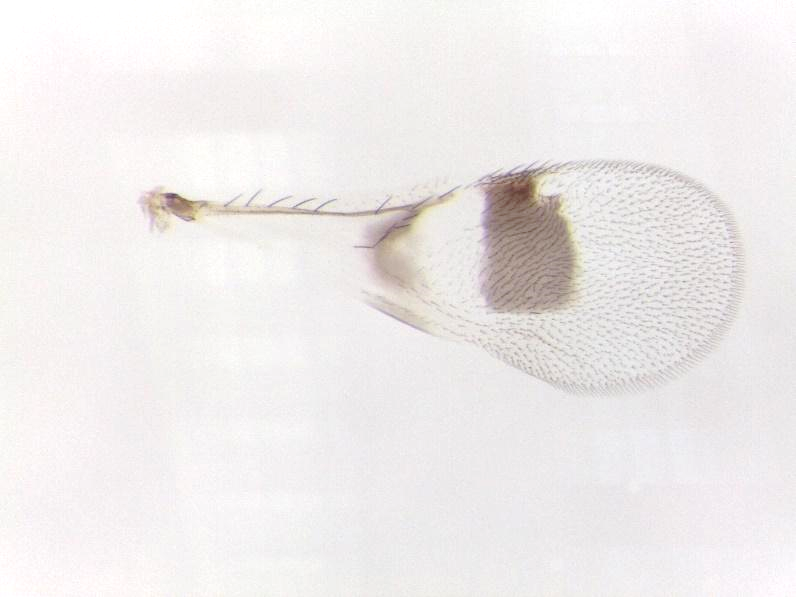

**Species 20. *Sycophila texana***

**Host gall: *Andricus quercuspetiolicola, Callirhytis seminator, Disholcaspis quercusglobulus* and unknown integral gall**

**Host plant: *Quercus alba, Q. bicolor*, and *Q. stellata***

**Locality: IA and TX**

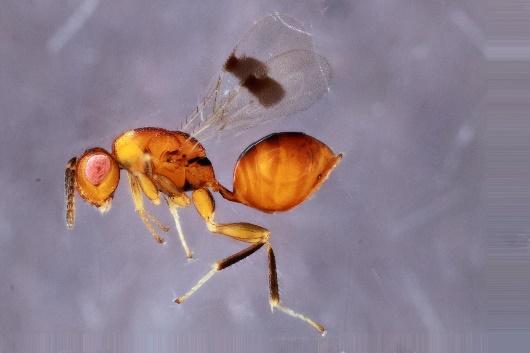

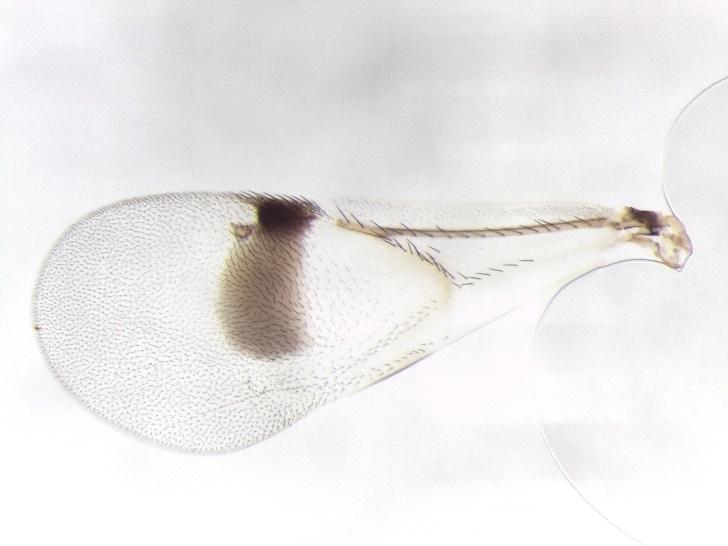

**Species 21. *Sycophila* sp6**

**Host gall *Callirhytis quercuspunctata*, host plant *Quercus palustris*, Locality: MO.**

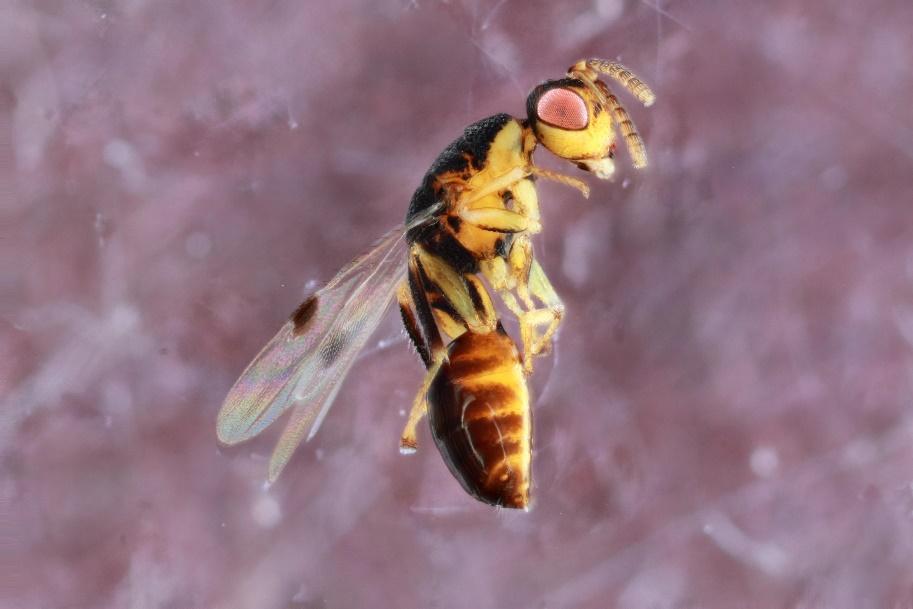

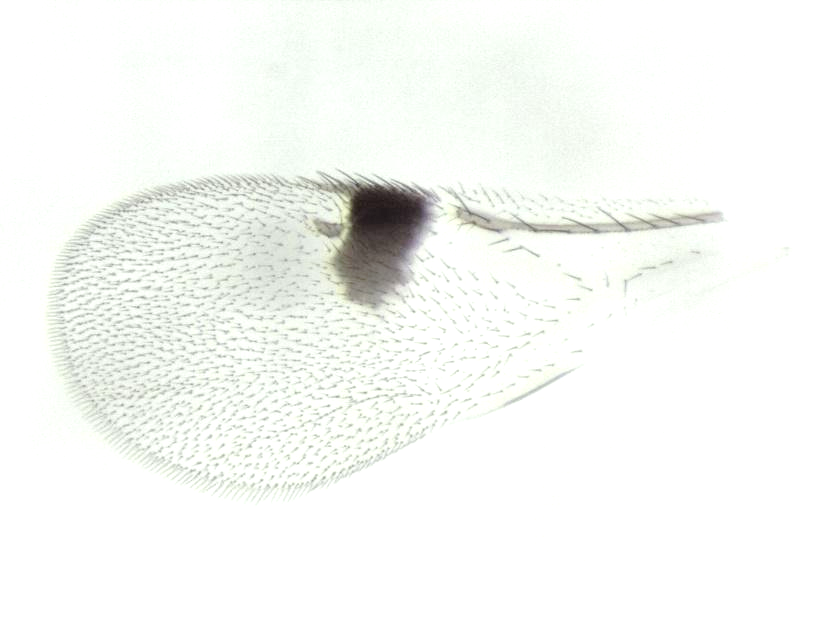

**Species 22. *Sycophila* nr. *nubilistigma***

**Host gall: *Amphibolips globus*, *Dryocosmus cinerceae*, and *D. quercuspalustris***

**Host plant: *Quercus palustris*, *Q. rubra* and *Q. myrtifolia***

**Locality: IA and FL.**

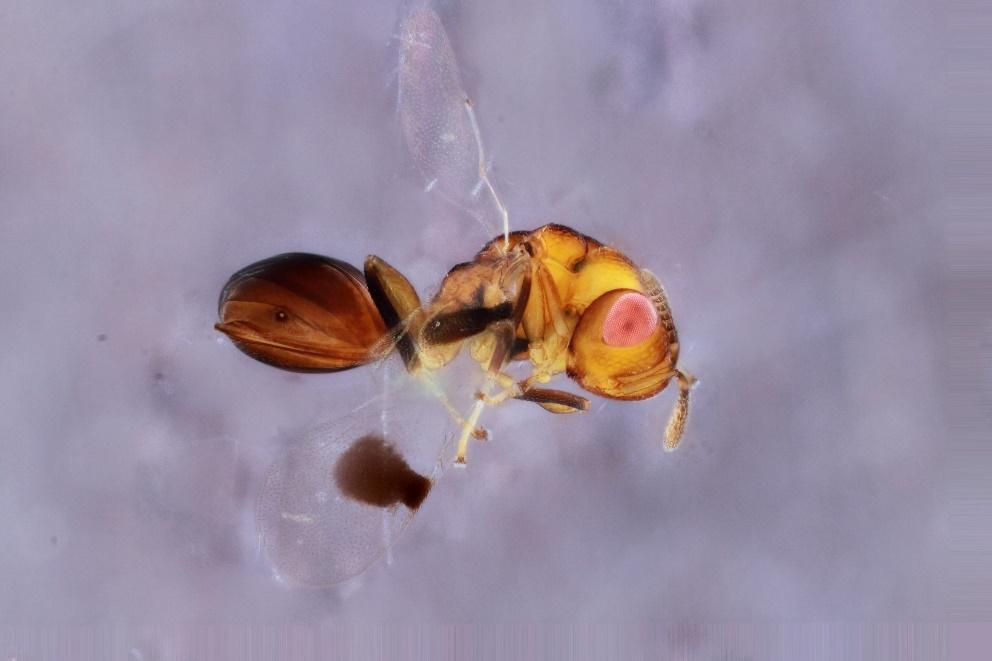

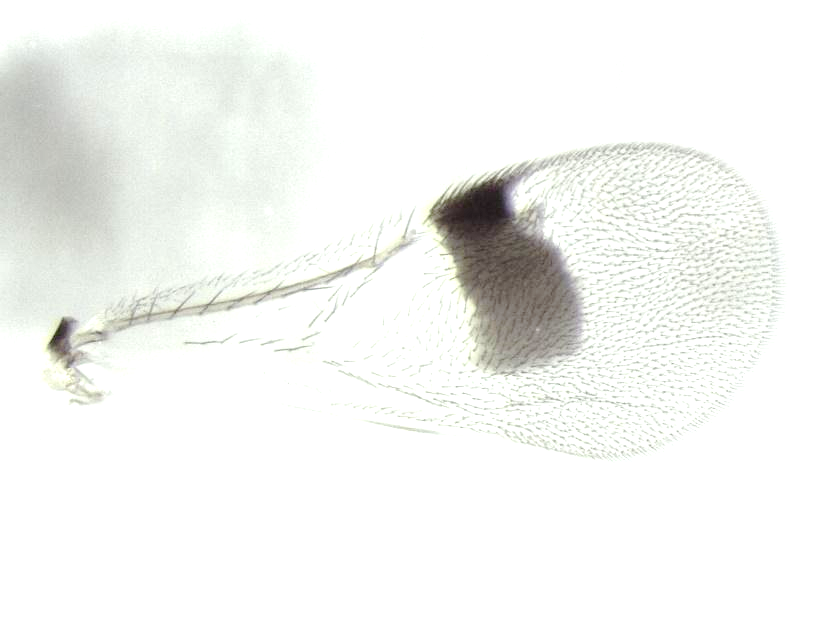

**Species 23. *Sycophila* nr. *globuli* (see Smith-Freedman et al. 2019 Supplemental Data for images)**

**Host gall: *Zapatella davisae***

**Host plant: *Quercus velutina***

**Locality: MA**

**Species 24. *Sycophila* nr. *dubia/globuli***

**Host gall: *Callirhytis quercusoperator*, *Cynips douglasii*, and *Neuroterus pallidus***

**Host plant: *Quercus bicolor*, *Q. lobata*, and *Q. velutina***

**Locality: CA and IA**

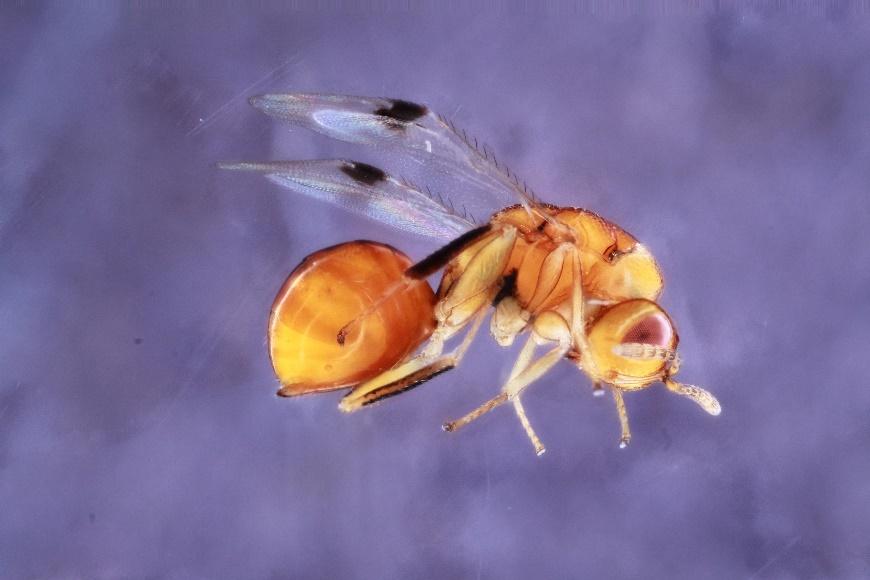

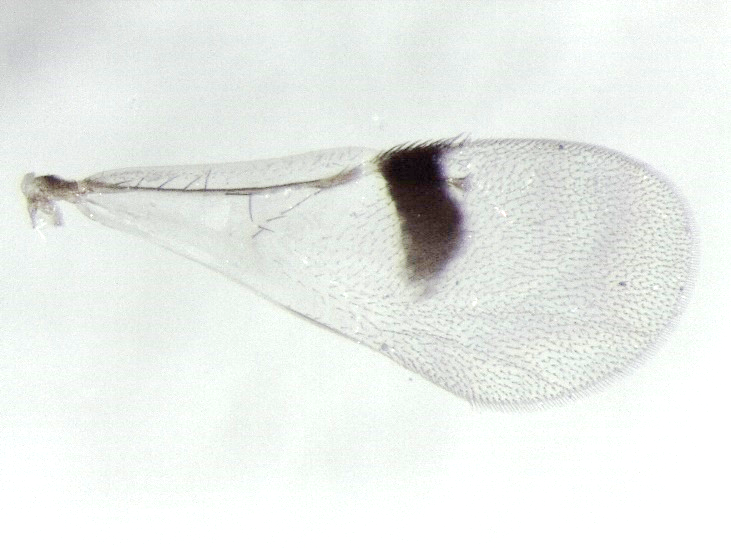

**Species 25. *Sycophila* sp7**

**Host gall: *Belonocnema kinseyi***

**Host plant: *Quercus fusiformis***

**Locality: TX**

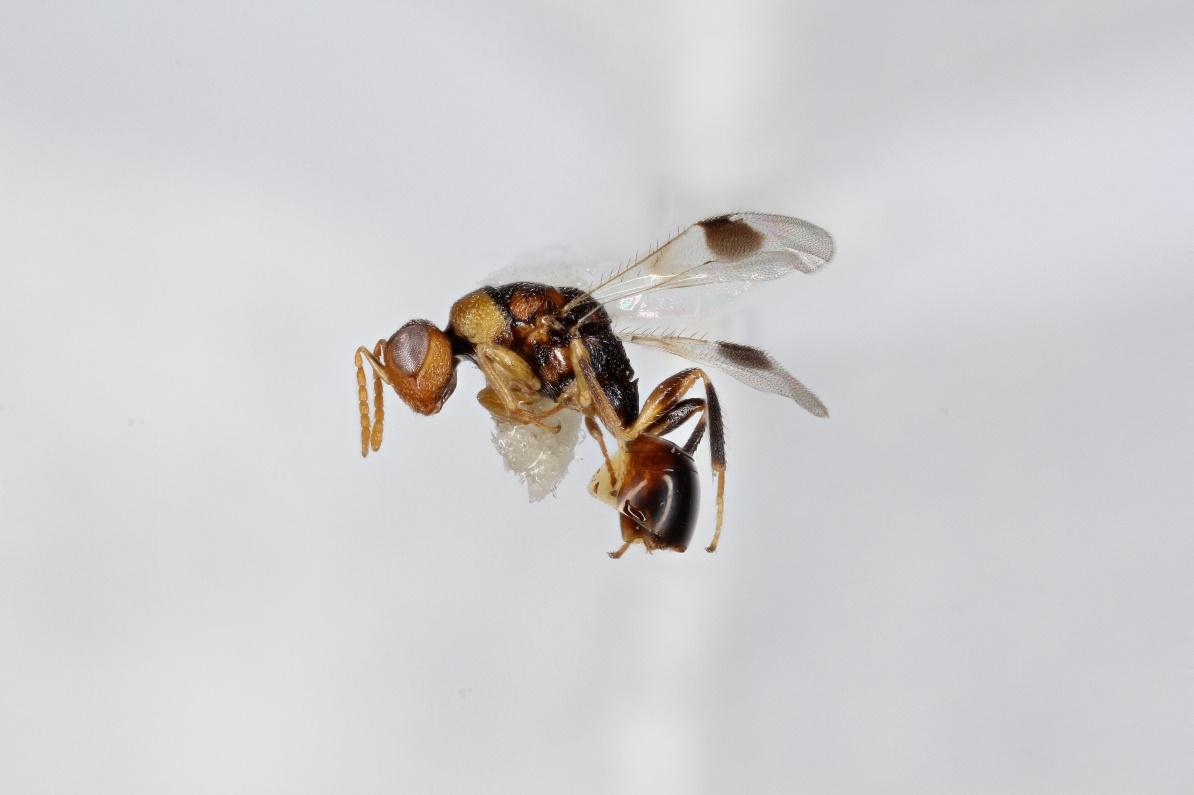

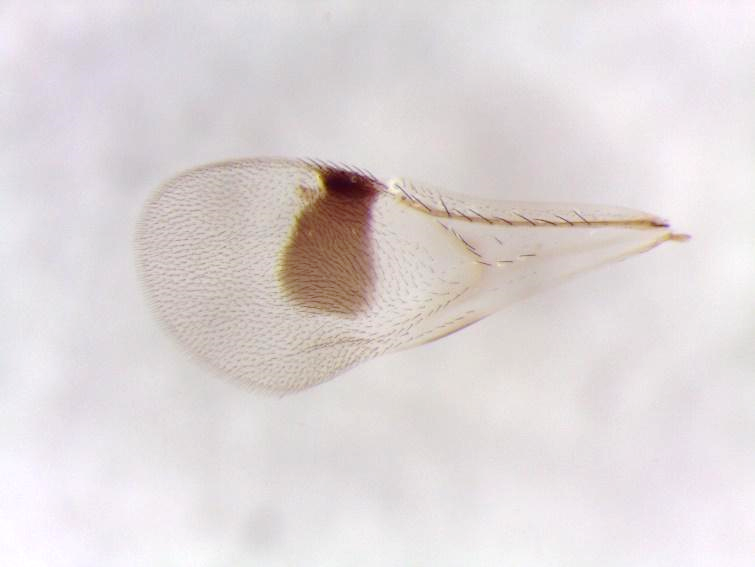

**Species 26. *Sycophila* nr. *lobatae***

**Host gall: *Callirhytis perdens***

**Host plant: *Quercus wislizeni***

**Locality: CA**

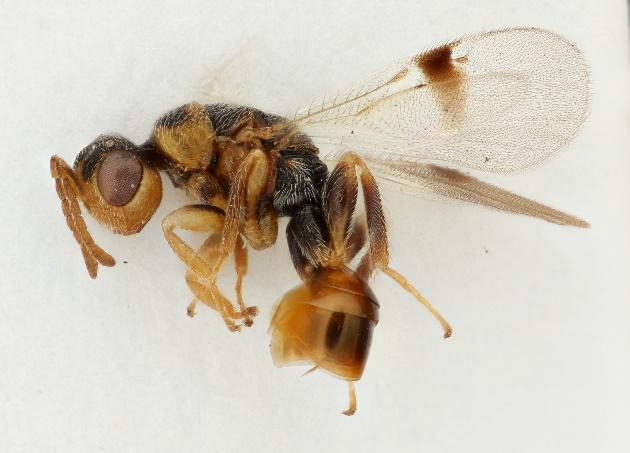

**Species 27. *Sycophila dubia*-1**

**Host gall: *Callirhytis quercuscornigera***

**Host plant: *Quercus palustris***

**Locality: MO**

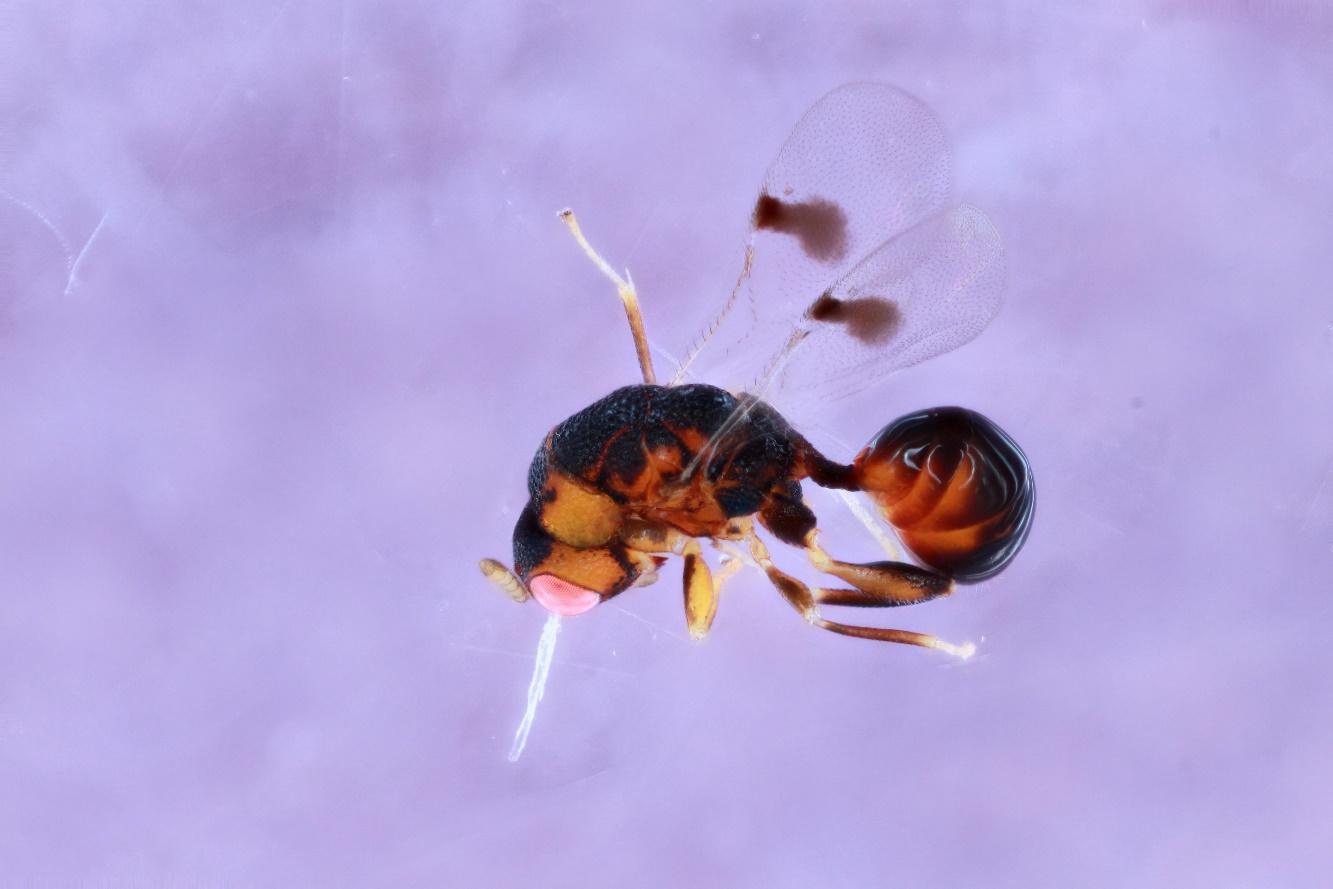

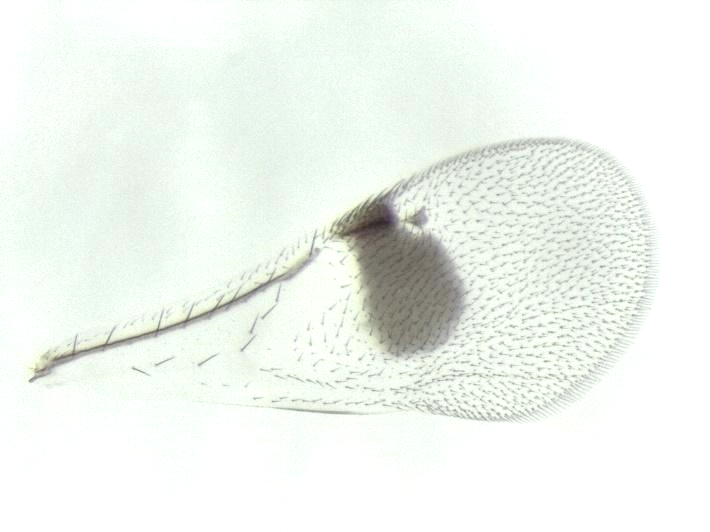

**Species 28. *Sycophila dubia*-2**

**Host gall: *Callirhytis quercusgemmaria***

**Host plant: *Quercus palustris***

**Locality: MO**

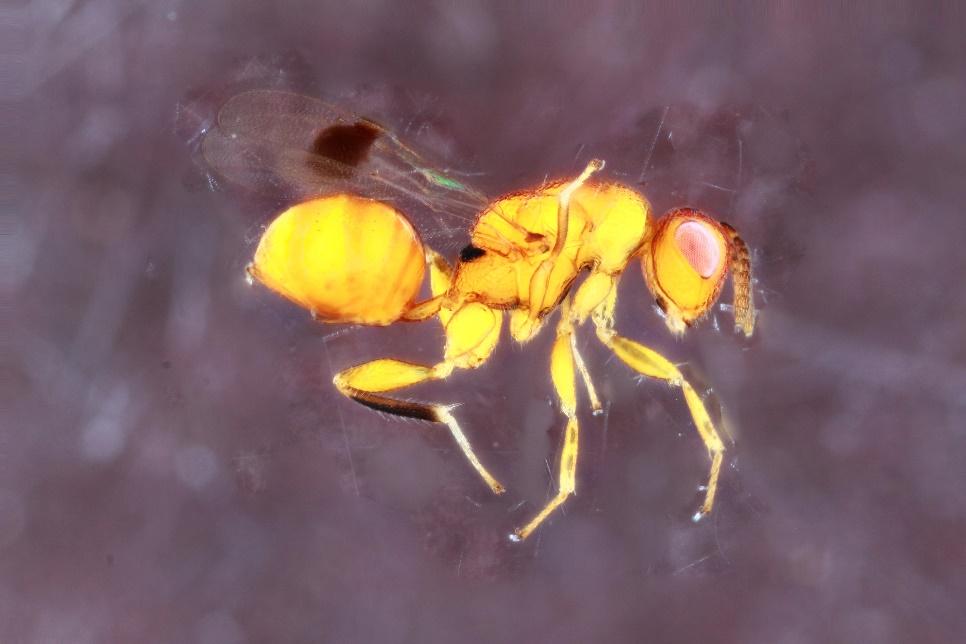

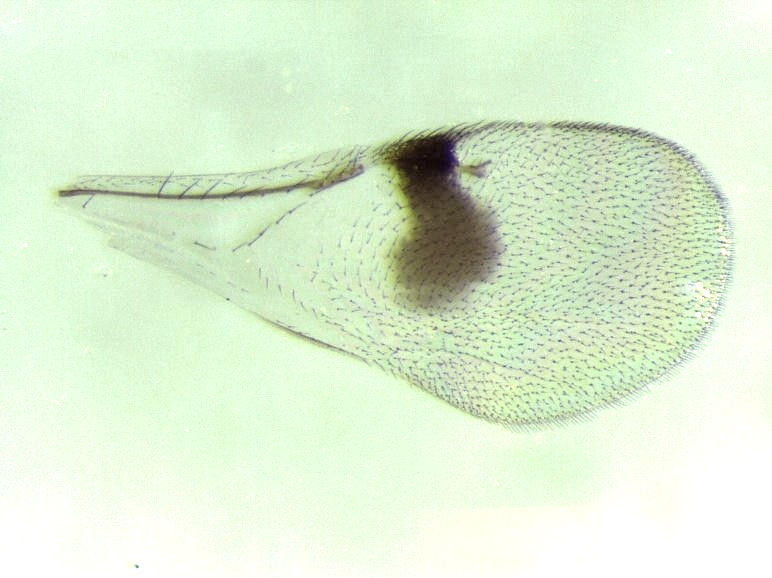

**Species 29. *Sycophila* nr. *nigriceps*-1**

**Host gall: *Melikaiella ostensackeni***

**Host plant: *Quercus rubra***

**Locality: IA**

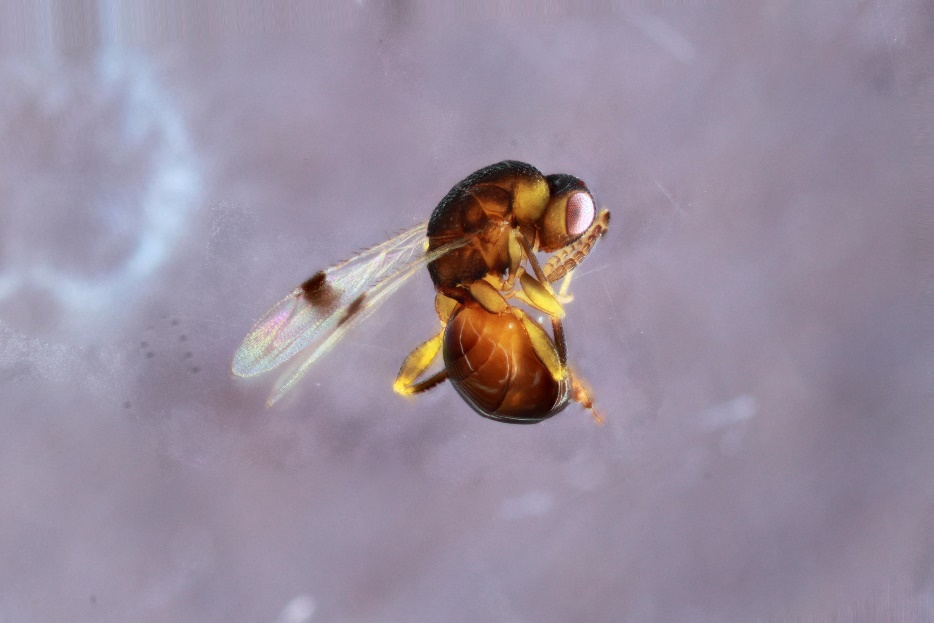

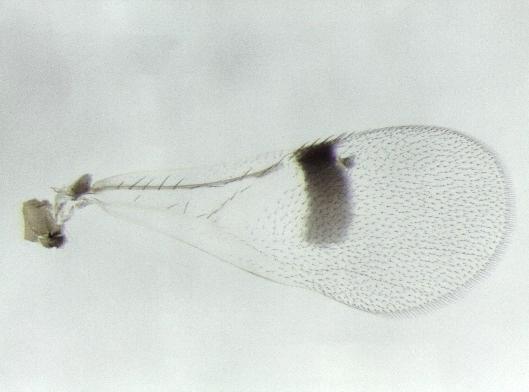

**Species 30. *Sycophila* nr. *nigriceps*-2**

**Host gall: *Disholcaspis quercusmamma***

**Host plant: *Quercus macrocarpa***

**Locality: IA**

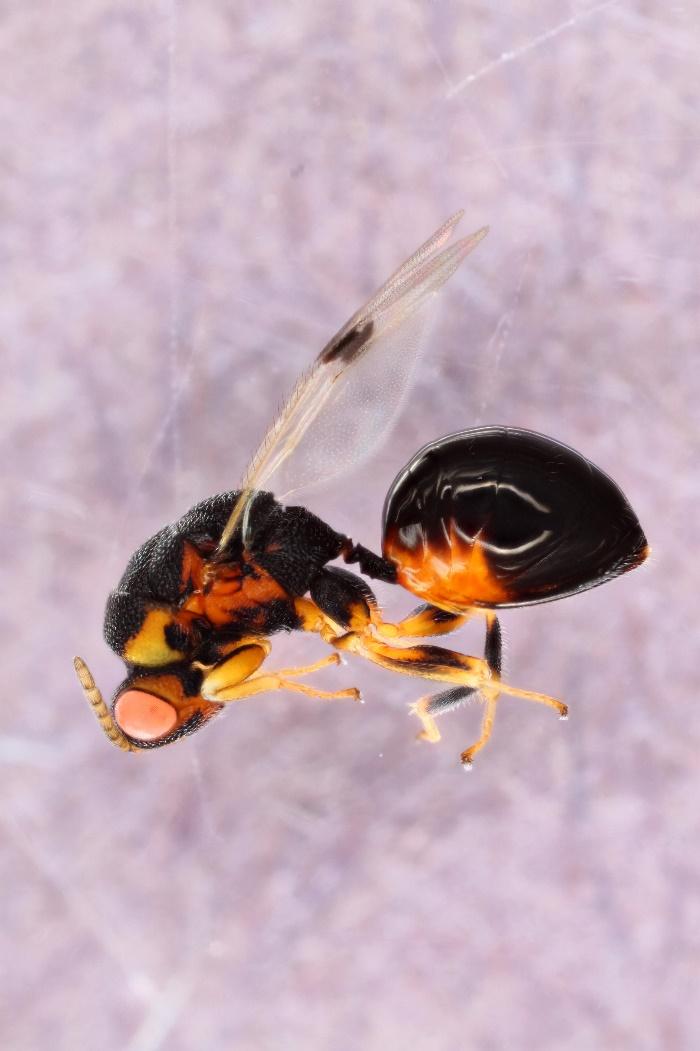

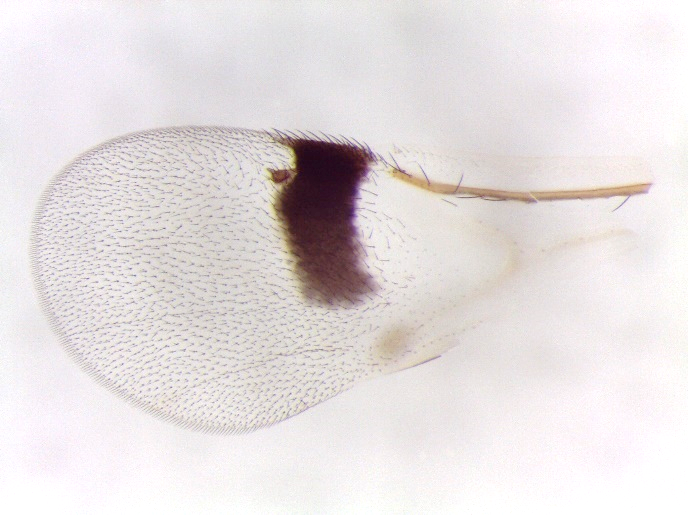

**Species 31. *Sycophila* nr. *occidentalis***

**Host gall: *Burnettweldia washingtonensis* and *Disholcaspis mellifica***

**Host plant: *Quercus garryana***

**Locality: BC and WA**

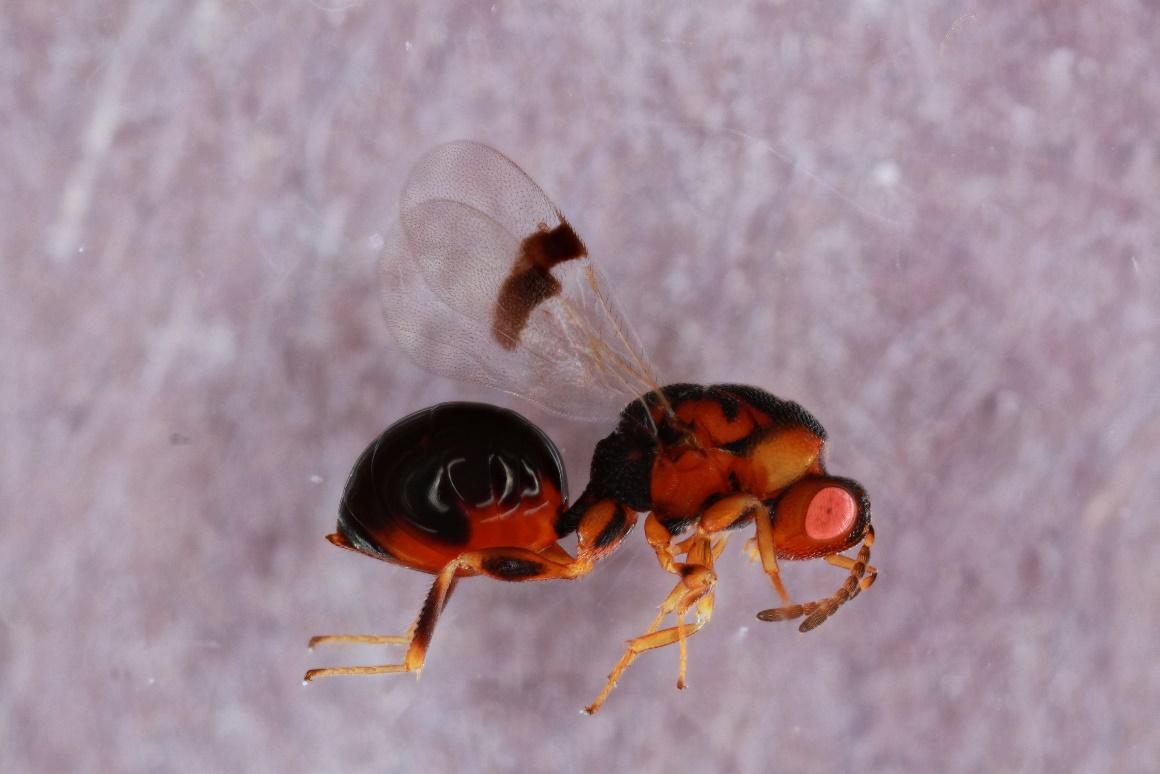

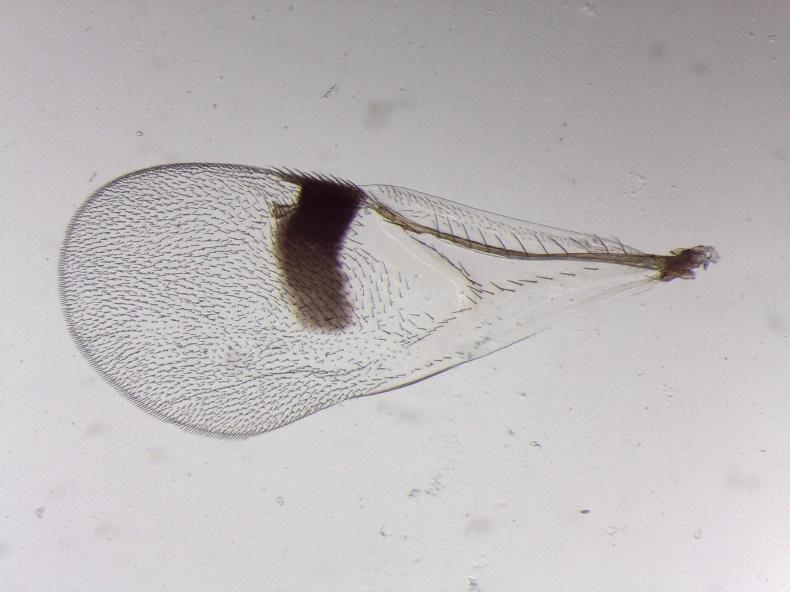

**Species 32. *Sycophila* sp8**

**Host gall: *Disholcaspis rubens***

**Host plant: *Quercus gambelii***

**Locality: AZ**

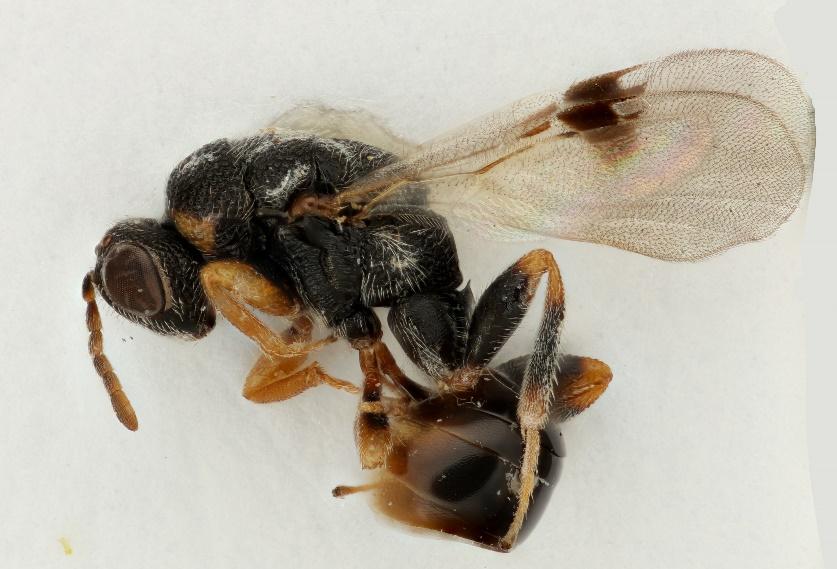

**Species 33. *Sycophila globuli***

**Host gall: *Disholcaspis quercusglobulus* and *D. quercusmamma***

**Host plant: *Quercus alba, Q. grisea*, and *Q. stellata***

**Locality: AZ, CO, and MO**

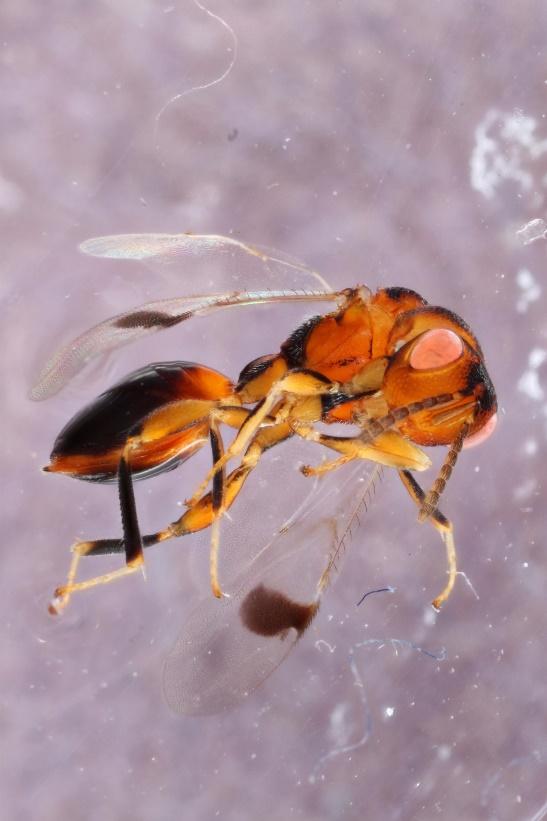

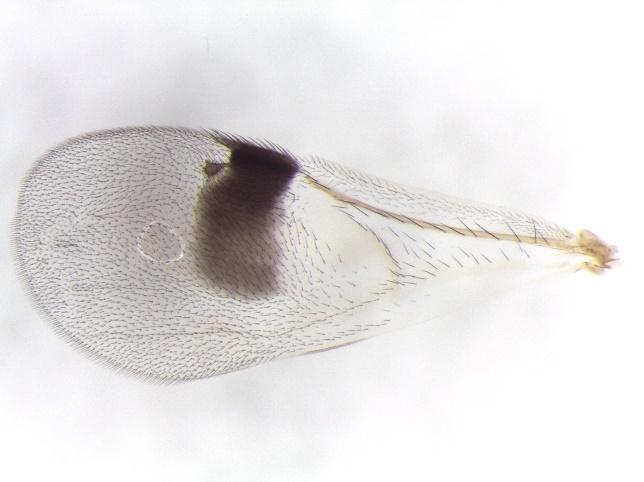

**Species 34. *Sycophila* nr. *wiltzae***

**Host gall: *Cynips douglasii***

**Host plant: *Quercus lobata***

**Locality: CA**

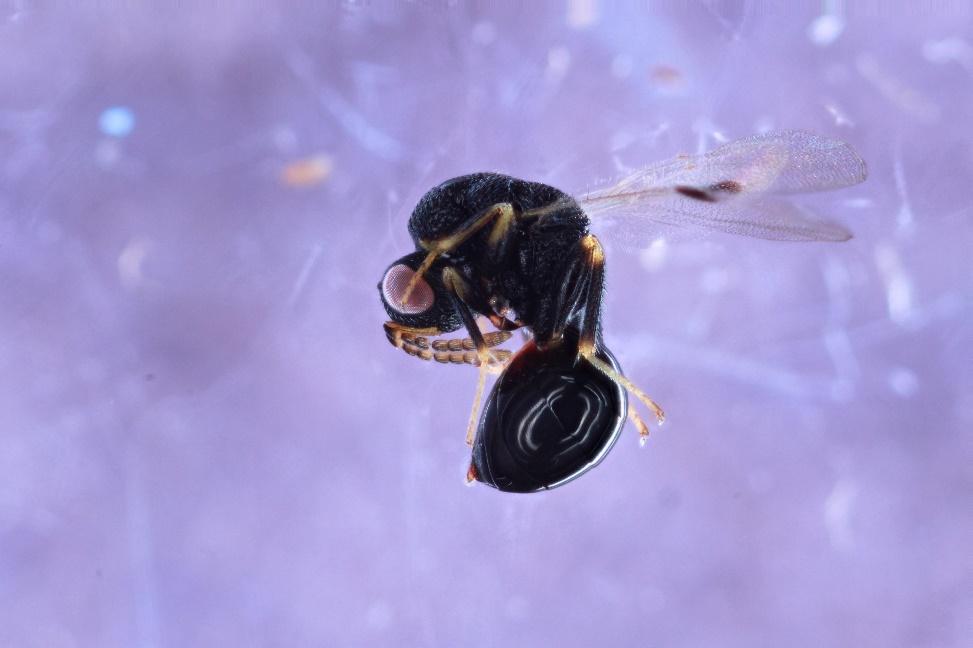

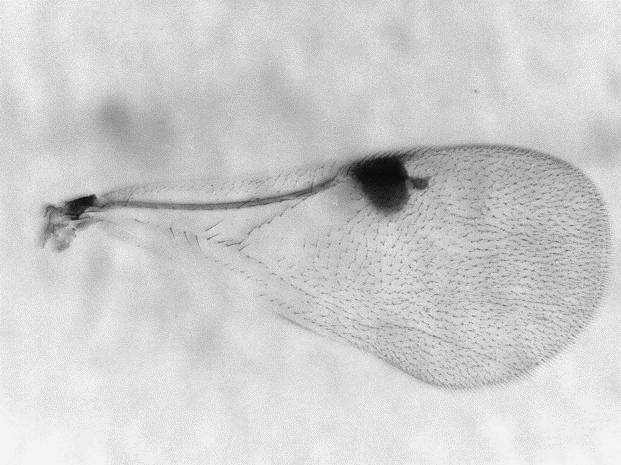

**Species 35. *Sycophila* sp9**

**Host gall: *Andricus incertus***

**Host plant: *Quercus bicolor***

**Locality: IA**

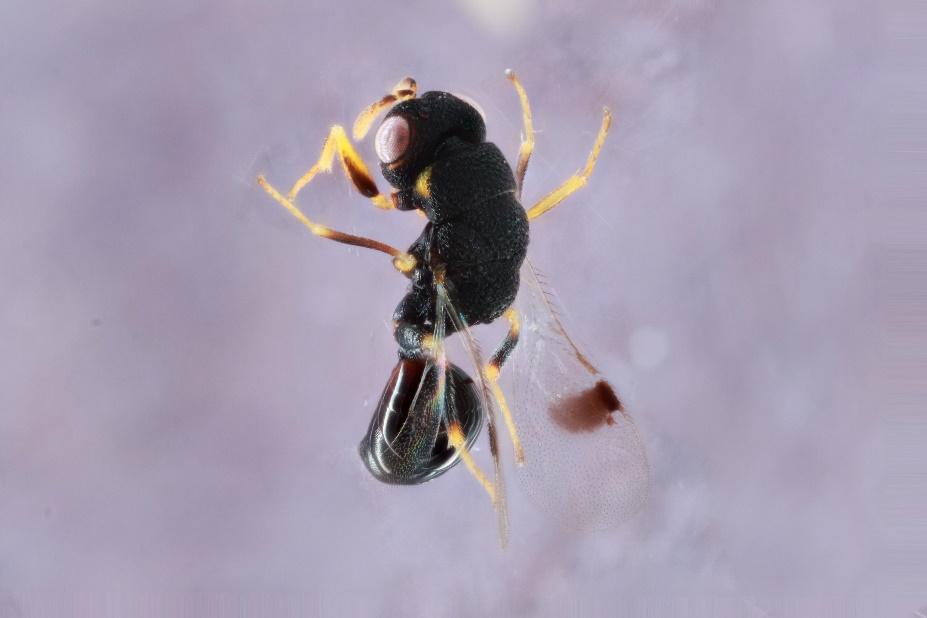

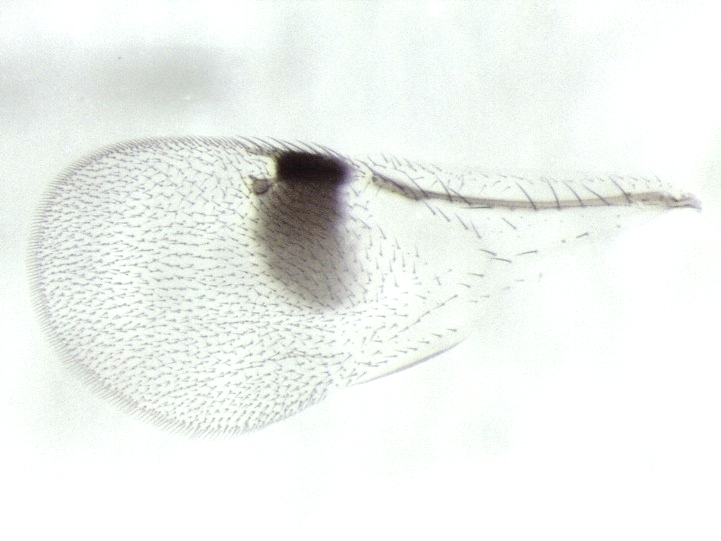
