## Supplementary figures and images for "Delimiting the cryptic diversity and host preferences of *Sycophila* parasitoid wasps associated with oak galls using phylogenomic data"

### Fig. S2

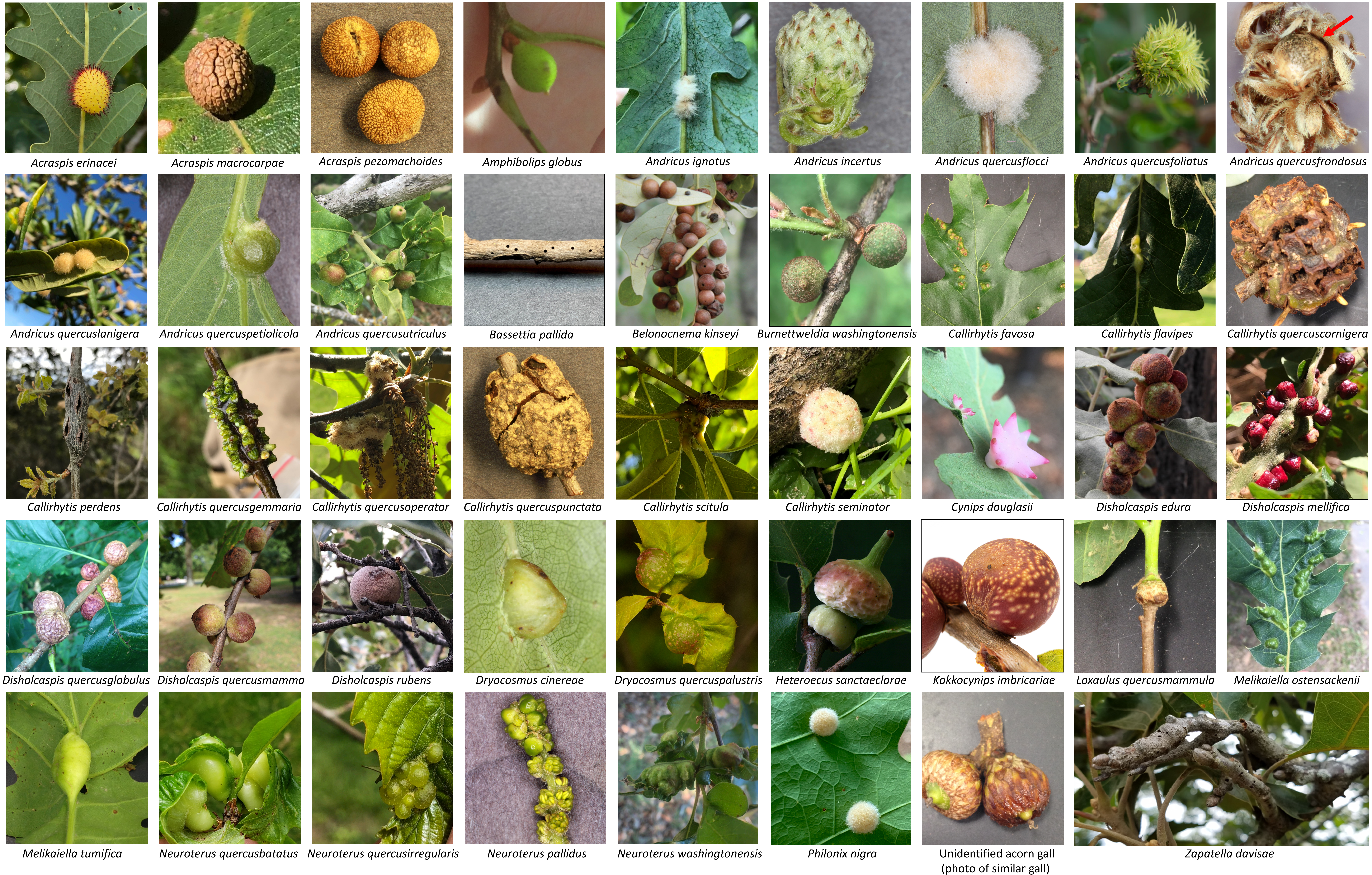

### Fig. S3

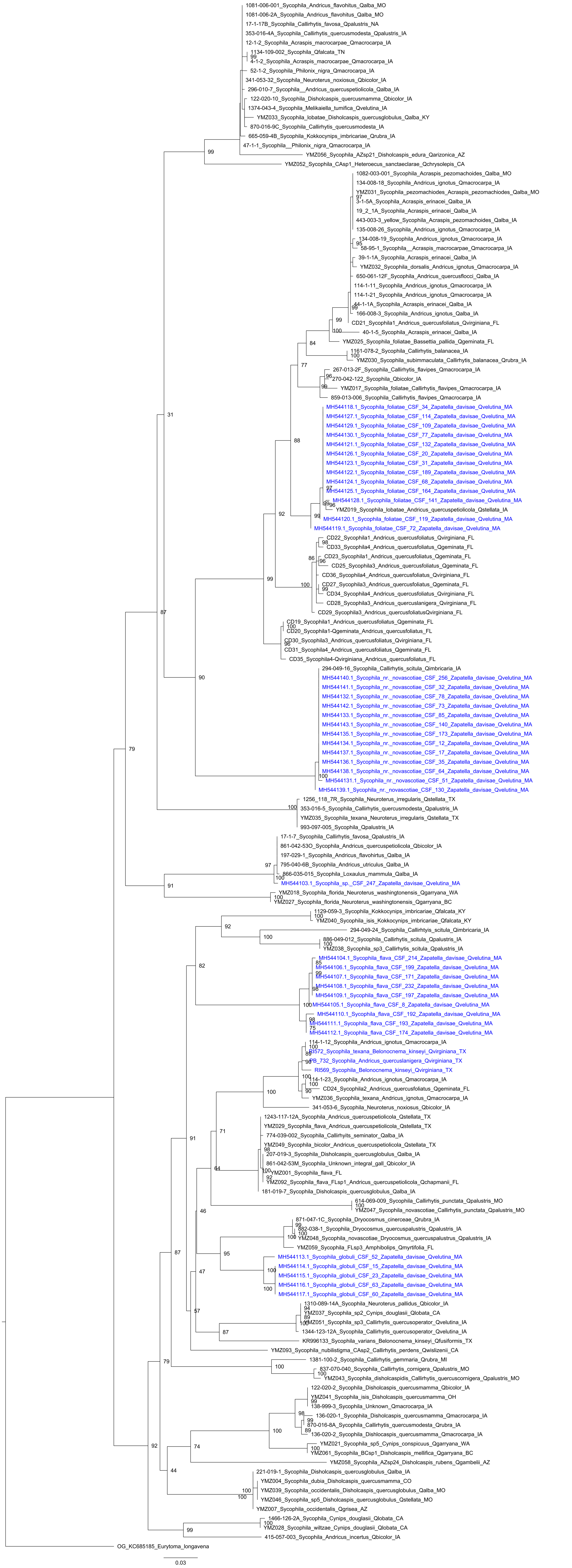

### Fig. S4

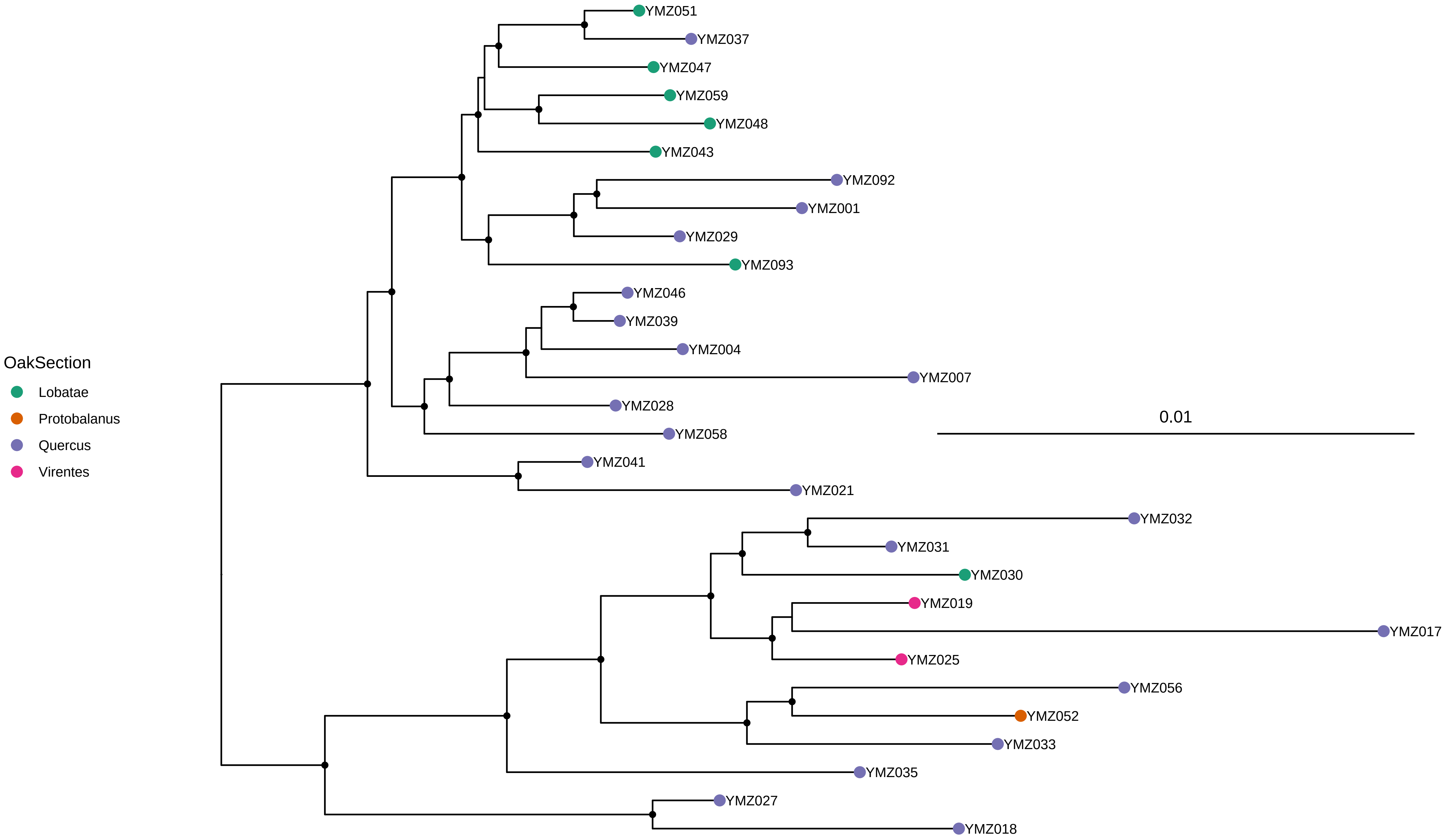

### Fig. S5

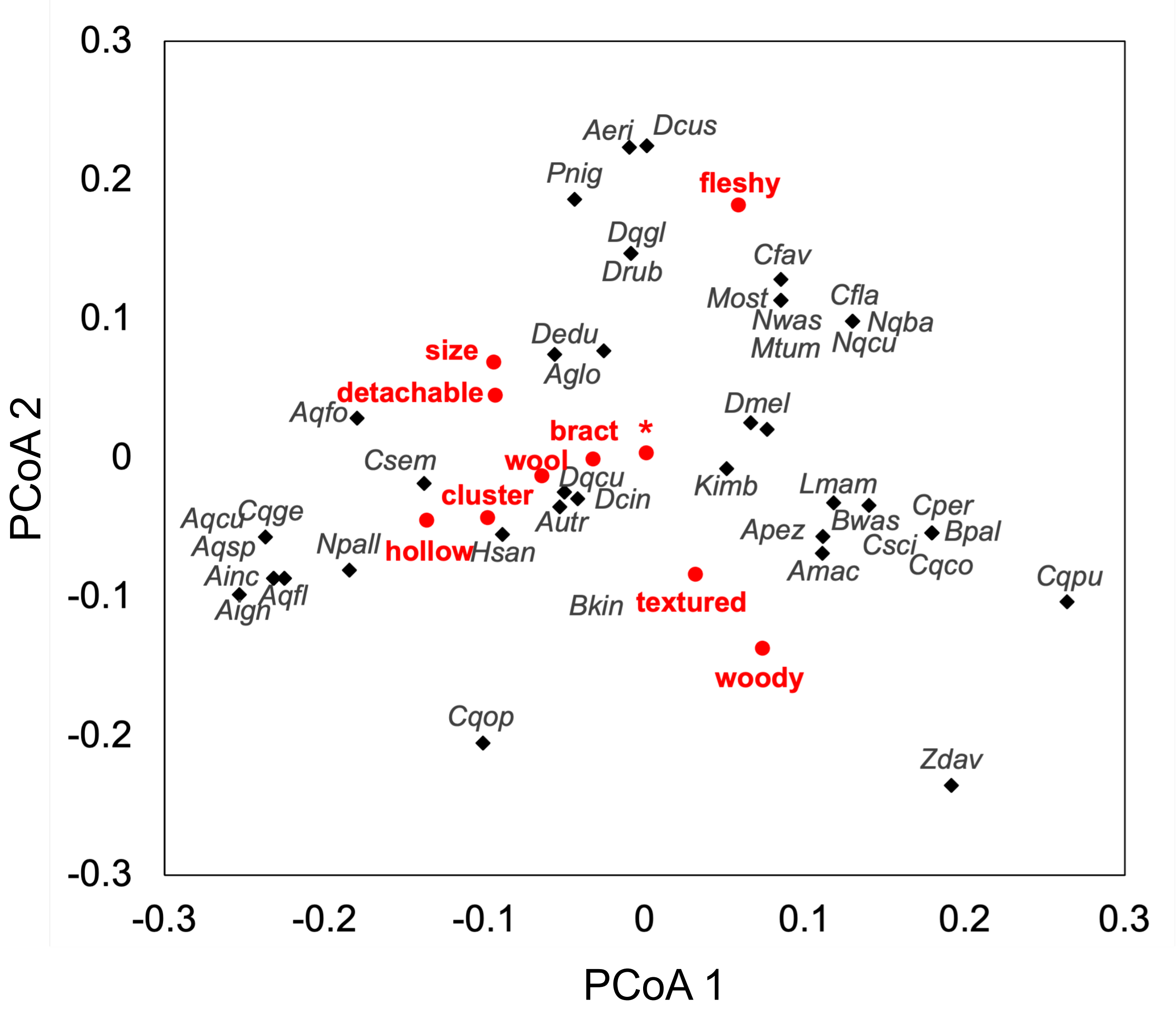

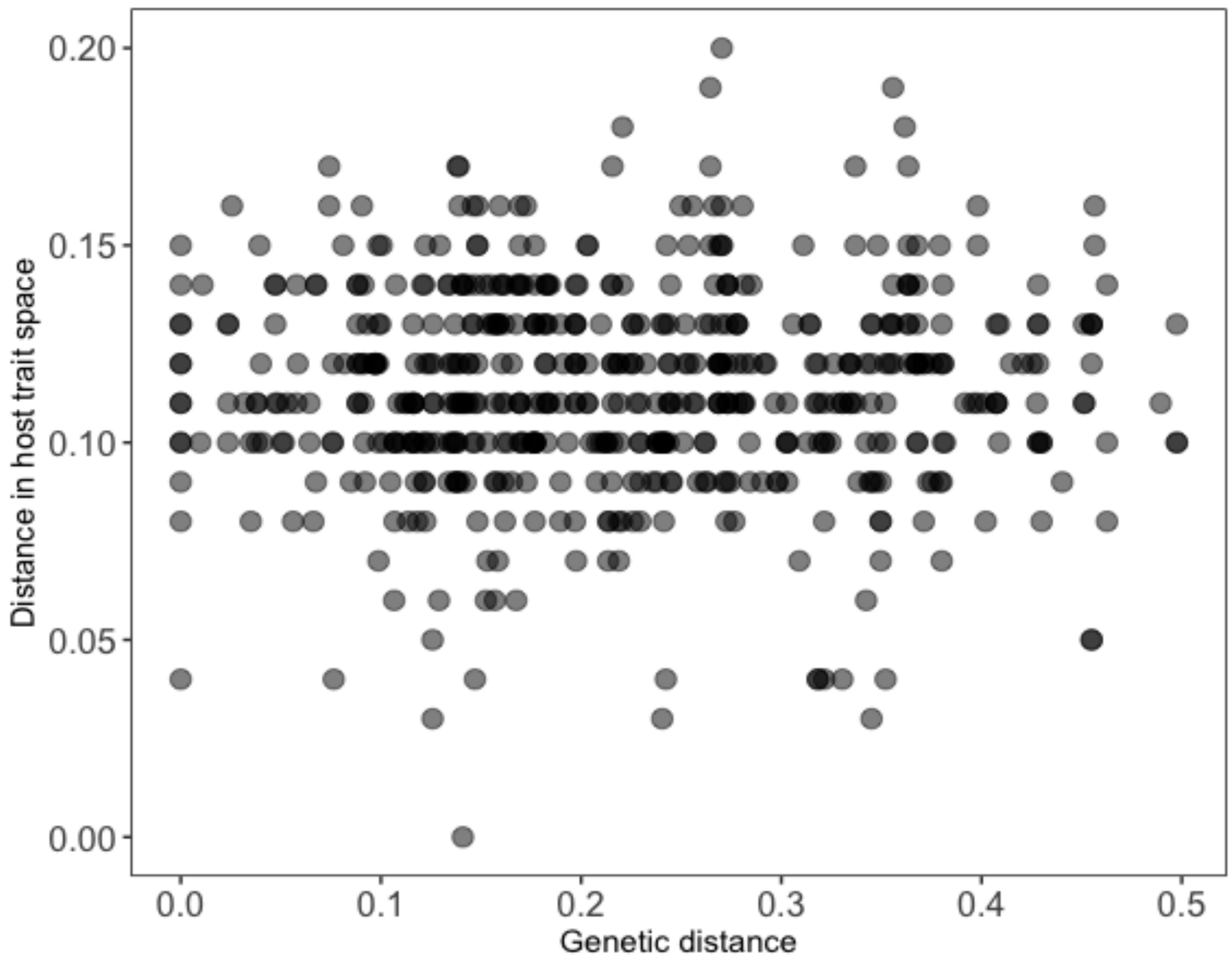
